# Rule-based and stimulus-based cues bias auditory decisions via different computational and physiological mechanisms

**DOI:** 10.1101/2021.12.09.471952

**Authors:** Nathan Tardiff, Lalitta Suriya-Arunroj, Yale E. Cohen, Joshua I. Gold

## Abstract

Expectations, such as those arising from either learned rules or recent stimulus regularities, can bias subsequent auditory perception in diverse ways. However, it is not well understood if and how these diverse effects depend on the source of the expectations. Further, it is unknown whether different sources of bias use the same or different computational and physiological mechanisms. We examined how rule-based and stimulus-based expectations influenced behavior and pupil-linked arousal, a marker of certain forms of expectation-based processing, of human subjects performing an auditory frequency-discrimination task. Rule-based cues consistently biased choices and response times (RTs) toward the more-probable stimulus. In contrast, stimulus-based cues had a complex combination of effects, including choice and RT biases toward and away from the frequency of recently presented stimuli. These different behavioral patterns also had: 1) distinct computational signatures, including different modulations of key components of a novel form of a drift-diffusion decision model and 2) distinct physiological signatures, including substantial bias-dependent modulations of pupil size in response to rule-based but not stimulus-based cues. These results imply that different sources of expectations can modulate auditory processing via distinct mechanisms: one that uses arousal-linked, rule-based information and another that uses arousal-independent, stimulus-based information to bias the speed and accuracy of auditory perceptual decisions.

**Author summary:** Prior information about upcoming stimuli can bias our perception of those stimuli. Whether different sources of prior information bias perception in similar or distinct ways is not well understood. We compared the influence of two kinds of prior information on tone-frequency discrimination: rule-based cues, in the form of explicit information about the most-likely identity of the upcoming tone; and stimulus-based cues, in the form of sequences of tones presented before the to-be-discriminated tone. Although both types of prior information biased auditory decision- making, they demonstrated distinct behavioral, computational, and physiological signatures. Our results suggest that the brain processes prior information in a form-specific manner rather than utilizing a general-purpose prior. Such form-specific processing has implications for understanding decision biases real-world contexts, in which prior information comes from many different sources and modalities.

## Introduction

The auditory system is sensitive to expectations [1–3]. Some expectations are instructed explicitly, such as through cues that carry information about the probability of occurrence of subsequent auditory stimuli. Other expectations are inferred explicitly or implicitly from the features of an auditory stimulus, including its temporal and structural regularities. Despite the importance of these “rule-based” and “stimulus-based” expectations, respectively, we lack a basic understanding about the effects of both forms of expectations on auditory perception, including whether they use distinct or shared computational and physiological mechanisms.

For example, how do rule-based and stimulus-based expectations affect categorical auditory decisions? In general, for non-auditory decision tasks, rule-based expectations can lead to biases in behavioral choices and response times (RTs) that are consistent with normative principles, including a tendency to make faster and more-prevalent choices to more-probable alternatives [4–7]. These effects are thought to be mediated via top-down input from higher-order brain regions to earlier sensory regions [8]. However, the role of such rule-based expectations in auditory decision-making is relatively unexplored (except see [1, 9]).

In contrast, stimulus-based expectations have been studied extensively and are thought to reflect, in part, bottom-up mechanisms in the auditory system, including neural adaptation to stimulus regularities [3,10–14]. In general, these adaptation-like mechanisms can have the opposite effect as rule-based cues, including potentiating responses to violations of stimulus regularities and biasing perception away from recent stimuli [15–18]. For example, after hearing a tone rising or falling in frequency, human subjects are more likely to judge that a subsequent tone is changing frequency in the opposite direction (e.g., falling after hearing a rising tone; [18]). However, it is also feasible that stimulus-based cues could be used to rapidly update prior beliefs, which could then bias subsequent decisions through similar top-down mechanisms used for rule-based effects [2,11,19–25].

We thus sought to answer a basic question about how the brain uses rule-based and stimulus-based information to form auditory decisions: Do decision biases elicited by explicit top-down rules versus those elicited by stimulus regularities use the same or different computational and physiological mechanisms? To answer this question, we recruited human subjects to perform a two-alternative, forced-choice frequency-discrimination task. These subjects reported whether a test stimulus (a tone burst that was embedded in a noisy background) was “low” frequency or “high” frequency. We manipulated both the signal-to-noise ratio (SNR) of the test tone (relative to the background) and two types of expectation-generating cues: 1) rule-based cues, in the form of visual stimuli indicating the probability of each test tone; and 2) stimulus-based cues, in the form of temporal sequences of tone bursts, akin to those used to study stimulus-specific adaptation and mismatch negativity [10,26,27], that immediately preceded the test tone. We also measured the subjects’ pupil diameter, which is a physiological marker of arousal that is sensitive to certain cognitive processes related to decision-making [28], including decision biases [9,21,29–31] and violations of top-down, but not bottom-up, expectations [32]. We found that rule-based and stimulus-based behavioral biases exhibited distinct computational and physiological signatures (as assessed via modeling and pupillometry, respectively) that may reflect differential contributions of top-down and bottom-up forms of expectation-dependent information processing in auditory perceptual decision-making.

## Results

Fifty human subjects performed a frequency-discrimination task in which they indicated with a button press whether a test tone was high frequency or low frequency (Figure 1). We titrated task difficulty by embedding the test tone in a background of white noise with different signal-to-noise ratios (SNRs). We manipulated prior information in a blockwise fashion for each subject. In the rule-based condition, prior to stimulus onset (Figure 1), one of three visual cues (randomly interleaved in mini-blocks of 48 trials) indicated the probability that the upcoming test tone would be low frequency or high frequency (cue high:low ratio: “low” = 1:5, “neutral” = 1:1, “high” = 5:1). In the stimulus-based condition, prior information was manipulated by presenting a temporal sequence of tone bursts (using the maximum SNR; 2–14 tone bursts per trial) prior to the test tone. This “pre-test” sequence varied in terms of its pattern of low and high frequencies but, on average, was not predictive of the frequency of the test tone. We also presented mixed conditions in which subjects encountered both rule-based and stimulus-based cues on each trial.

**Figure 1.**
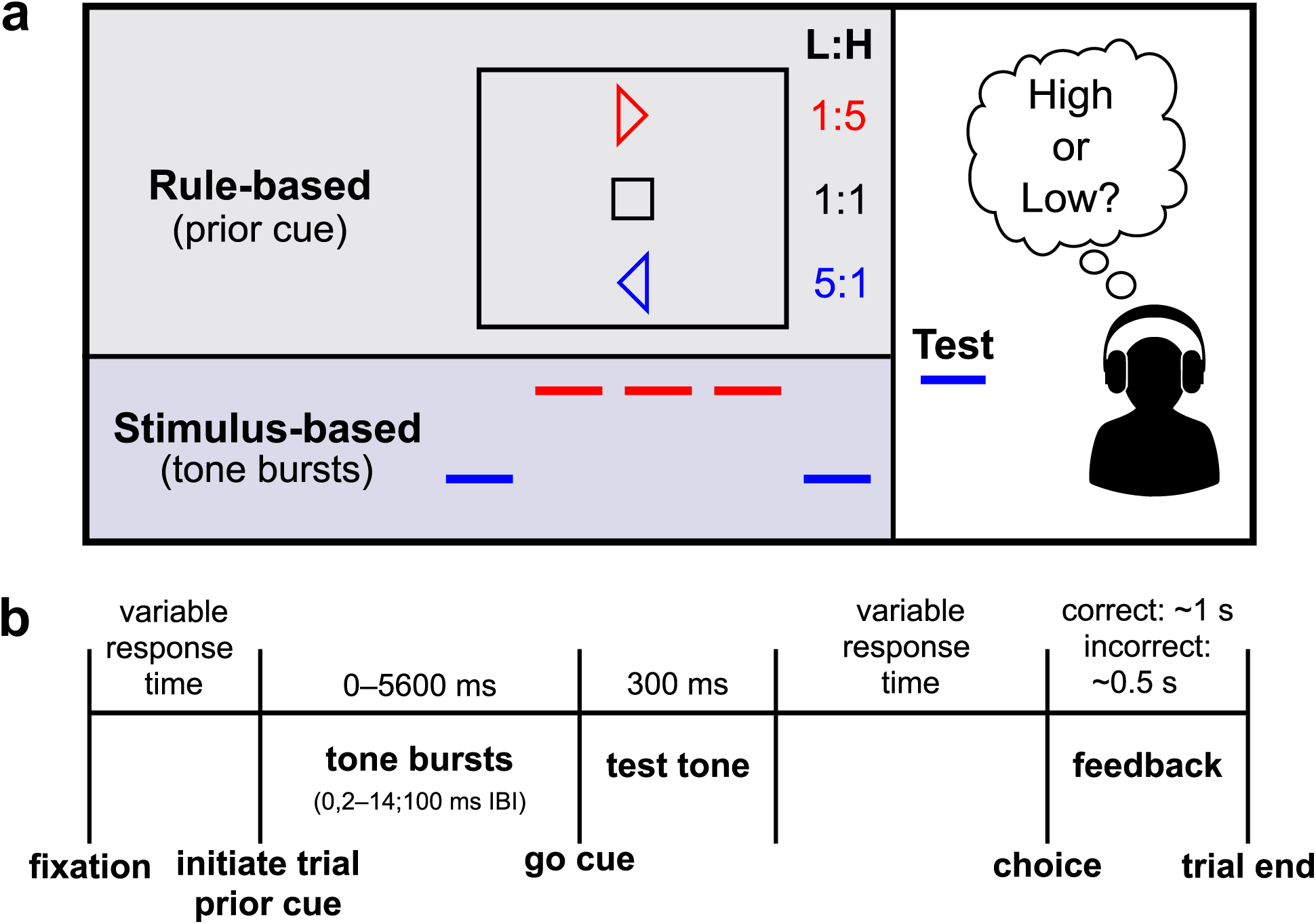
Task structure. **a** Subjects indicated with a button press whether a test tone was “high” frequency or “low” frequency. In the rule-based condition, the test tone was preceded by a visual stimulus (prior cue) that indicated the probability that the upcoming tone would be low (L) or high (H) frequency, with the probability determined by the L:H ratio. In the stimulus-based condition, the test tone was preceded by a variable-length, temporal “pre-test” sequence of low- and high- frequency tones. On average, this sequence was not predictive of the test tone. In the mixed conditions, subjects received both rule-based and stimulus-based cues on each trial. **b** Detailed trial timing. After attaining fixation, the subject initiated the trial with a button press. The subject was then able to report their decision with a second button press at any time following the “go cue” (up to 2000–3000 ms).

### Rule- and stimulus-based biases had different effects on choices and RTs

In the rule-based condition, the subjects’ choices and RTs showed consistent biases in accordance with the prior cues (Figure 2a). In particular, the subjects tended to choose the more-probable option more often and more quickly than a neutral or less-probable option. For example, when the prior cue indicated that the test tone was more likely to be high frequency, subjects chose high frequency more often and responded faster when the test tone was high frequency than when it was low frequency.

**Figure 2.**
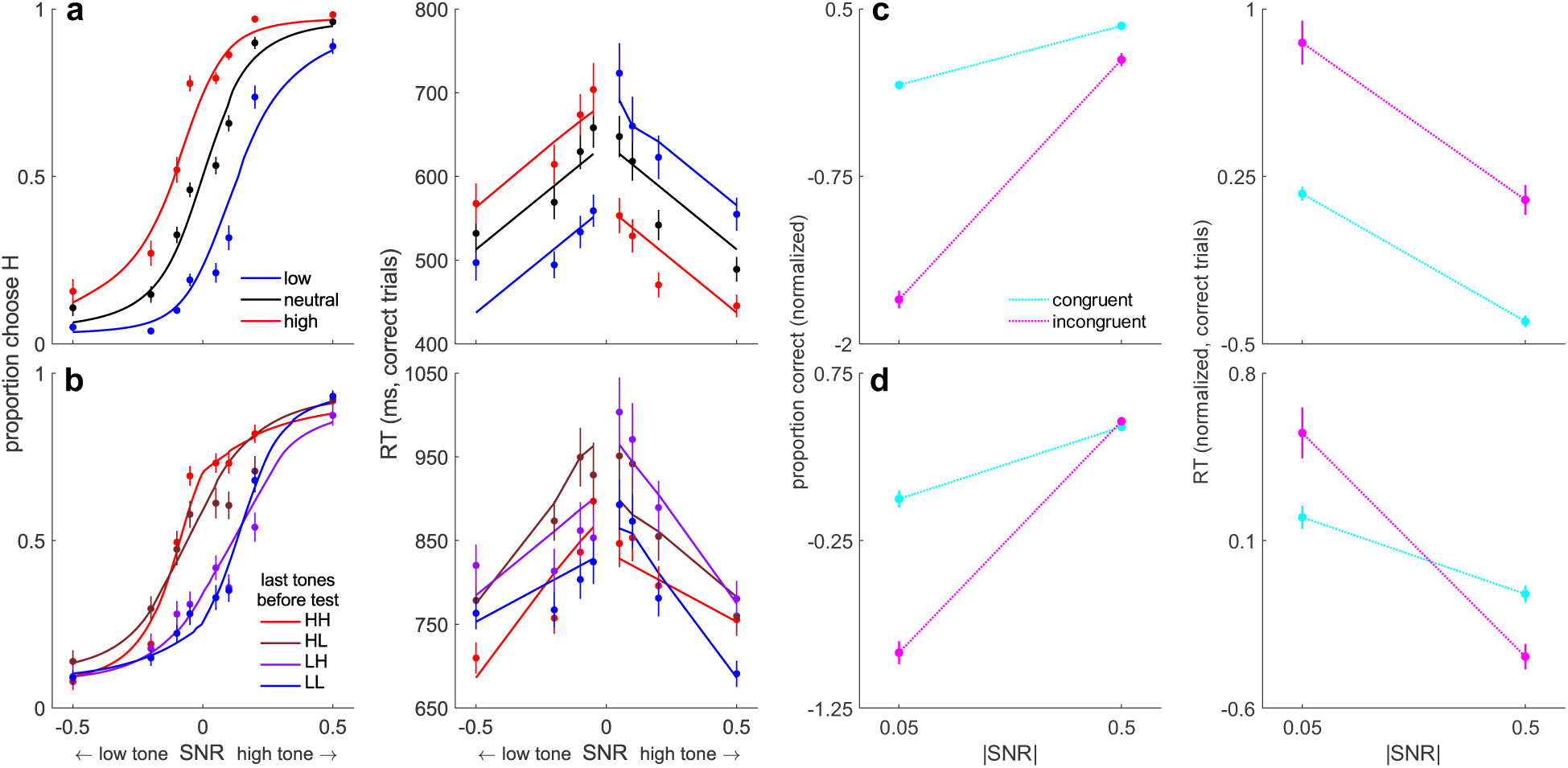
Behavioral summary. **a** Psychometric (left) and chronometric (right) functions from the rule-based condition for each prior cue (see legend). **b** Psychometric (left) and chronometric (right) functions from the stimulus-based condition for different patterns of the last two pre-test tones (see legend; HH = the last two tones were high, HL = the last tone was high and the second-to-last tone was low, etc.). By convention, negative and positive SNRs are low-frequency and high-frequency test tones, respectively. **c** Choice (left) and RT (right) interaction plots from the rule-based condition for trials in which the cue and test tone were “congruent” (e.g., a low cue followed by a low-frequency tone) or “incongruent” (see legend), at the lowest and highest unsigned SNRs. **d** Same as **c**, but for the stimulus-based condition. Here, only trials in which the last two pre-test tones were the same (HH and LL trials) were included in the analysis. In **a–d**, points are averages over subjects’ data, and error bars are standard errors of the mean (SEM). Within-subject RT averages in **a** and **b** were computed using medians, in accordance with the linear mixed-effects analysis. In **c** and **d**, choice and RT data were z-scored within subjects to facilitate comparisons. In **a** and **b**, solid lines are fits from the logistic (left) and linear (right) models for each condition (see text). Deviations of the RT fits from linearity are due to averaging predictions across subjects. In **c** and **d**, dotted lines are for visualization purposes only.

To quantify these effects, we first fit logistic models to choice data and linear models to median RTs from correct trials (Figure 2a; Tables 1–2). The choice bias was well characterized as an additive shift (bias) of the logistic psychometric function in favor of the prior (mean ΔAIC base versus bias = 54.60). These fits were only marginally improved by allowing the slope (sensitivity) of the psychometric function to vary with the prior (mean ΔAIC bias vs. bias + sensitivity = 0.29). A Bayesian random-effects analysis confirmed the bias-only model as the most likely model (protected exceedance probability = 0.82).

**Table 1.**
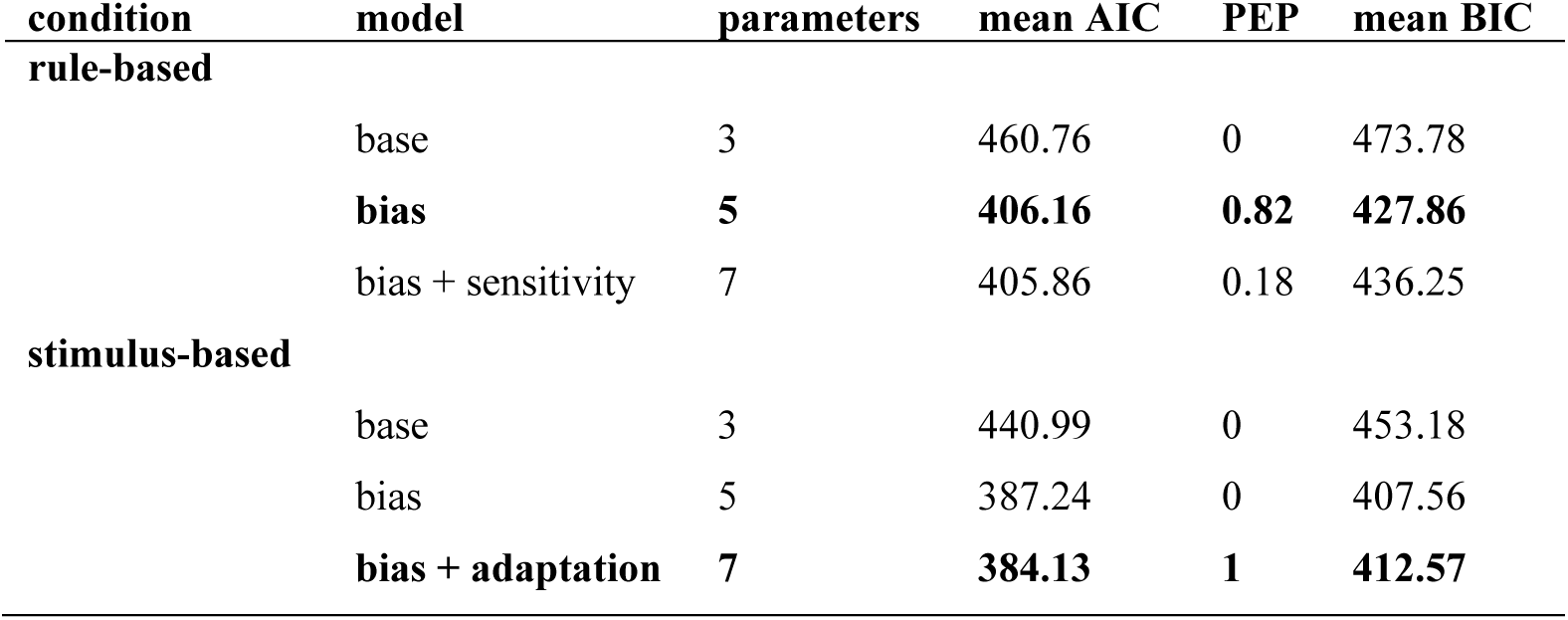
Logistic model-fit comparisons. AIC: Akaike information criterion; PEP: protected exceedance probability; BIC: Bayesian information criterion. We selected best-fitting models (bold) on the basis of AIC and PEP.

| condition | model | parameters | mean AIC | PEP | mean BIC |
| --- | --- | --- | --- | --- | --- |
| **rule-based** |  |  |  |  |  |
|  | base | 3 | 460.76 | 0 | 473.78 |
|  | **bias** | **5** | **406.16** | **0.82** | **427.86** |
|  | bias + sensitivity | 7 | 405.86 | 0.18 | 436.25 |
| **stimulus-based** |  |  |  |  |  |
|  | base | 3 | 440.99 | 0 | 453.18 |
|  | bias | 5 | 387.24 | 0 | 407.56 |
|  | **bias + adaptation** | **7** | **384.13** | **1** | **412.57** |

**Table 2.** Mixed-effects RT model ANOVAs. *NDF:* numerator degrees of freedom; *DDF:* denominator degrees of freedom. Denominator degrees of freedom were estimated using the Kenward-Roger method.

| condition | term | *F* | *NDF,DDF* | *p* |
| --- | --- | --- | --- | --- |
| **rule-based** |  |  |  |  |
|  | SNR | 75.57 | 1,48.02 | < 0.0001 |
|  | congruence | 160.43 | 2,65.08 | < 0.0001 |
| **stimulus-based** |  |  |  |  |
|  | SNR | 56.66 | 1,44.01 | < 0.0001 |
|  | congruence | 27.76 | 3,74.02 | < 0.0001 |
|  | congruence×SNR | 11.58 | 3,197.17 | < 0.0001 |

The RTs showed similar rule-based biases, along with a sharp discontinuity within the low- and high-prior conditions depending on whether subjects chose with (“congruent”; i.e., the prior and choice were toward the same frequency) or against (“incongruent”; i.e., the prior and choice were toward opposite frequencies) the cued prior (modulation by congruence, *F*(2,65.08) = 160.43, *p* < 0.0001; Table 2). For example, on average, RTs at the lowest SNR were >100-ms faster when

subjects chose high frequency with a high-prior cue or chose low frequency with a low-prior cue than when they made a choice that was incongruent with the prior (Figure 2a, right). Congruent RTs were faster than both incongruent RTs (*B =* 128.17, *t*(226.27) = 17.83 *p*_corrected_ < 0.0001) and RTs on neutral-prior trials (*B =* 75.48, *t*(51.03) = 9.19, *p*_corrected_ < 0.0001), whereas incongruent RTs were slower than neutral-prior RTs (*B = −*52.69, *t*(49.30) = −6.04, *p*_corrected_ < 0.0001). Thus, on average, the prior cues resulted in choices that were more common and faster to the more probable alternative.

In the stimulus-based condition, the subjects’ choices and RTs showed a more complex pattern. Figure 2b shows choices and RTs plotted as a function of the last two pre-test tones for a given trial. When the SNR of the test tone was low, choices were biased toward the frequency of the most-recently heard pre-test tones, and RTs were faster when the choice was congruent with the most-recently heard pre-test tones. These effects are evident in the lateral shifts in the psychometric function and discontinuities (vertical shifts) in the RTs at low SNRs. These effects are similar to those found in the rule-based condition (see Figure 2a).

In contrast, at higher test-tone SNRs, biases were in the opposite direction, such that subjects were, on average, more likely and faster (RT SNR×congruence interaction, *F*(3,197.17) = 11.58, *p* < 0.0001; Table 2) to respond with the alternative that was incongruent with the most-recently heard pre-test tones. That is, there was a frequency-specific adaptation-like effect that tended to weaken the perceptual impact of repeated, high-SNR stimuli. For example, subjects were less likely and slower to report high frequency for a high-SNR, high-frequency test tone after having heard two high-frequency tones than after having heard two low-frequency tones (Figure 2b, blue data points vs. red data points at SNR = 0.5; post-hoc test of RT interaction, *B =* 112.26, *t*(442.16) = 5.06, *p*_corrected_ < 0.0001).

Thus, there appeared to be a fundamental difference in the effects of rule-based versus stimulus-based expectations on choices and RTs, particularly at high test-tone SNRs. Specifically, for the rule-based condition, choices and RTs were consistently biased in the direction indicated by the cue. These effects were present at all test-tone SNRs but tended to be weaker, on average, when the test-tone SNR was high (which is consistent with normative perceptual decisions that show relatively weaker effects of priors when the sensory evidence is strong [33]; Figure 2c). In contrast, for the stimulus-based condition, choices and RTs tended to be biased in the direction of the pre-test tones when the test-tone SNR was low (akin to the bias in the rule-based condition) but in the opposite direction when the test-tone SNR was high (Figure 2d). The effect at high test-tone SNR resembles the effects of stimulus-specific adaptation in the auditory system, in which repeated tones of a specific frequency diminish neural responses to that tone frequency and potentiate responses to tones of a different (deviant) frequency [10,26,27,34]. To quantify the differences between conditions, we conducted an interaction analysis which asked how choice and RT varied along three dimensions: SNR (lowest versus highest), the congruence of the cue and the tone (congruent versus incongruent), and condition (rule-based versus stimulus-based). We found that both choice and RT data depended on a three-way SNR×congruence×condition interaction (choice: χ^2^(1) = 10.82, *p* = 0.001; RT: *F*(1,428.46) = 6.54, *p* = 0.01).

To further decompose the different effects of the pre-test tone sequence on choice and estimate the contributions of individual pre-test tones presented at different positions in the sequence relative to the (final) test tone, we fit a logistic model that included for each pre-test tone position: 1) an additive bias term describing the degree to which choice was biased in the same (positive values) or opposite (negative values) direction as the frequency of that pre-test tone; and 2) an SNR-dependent (unsigned SNR), adaptation-like term describing the change in discriminability attributable to that pre-test tone that would result in more (positive values) or fewer (negative values) choices to the frequency of that tone. These fits yielded, on average, a positive bias and a negative SNR-dependent adaptation-like component. Both the bias and adaptation-like terms had a strong recency effect in which the final 2–3 tones before the test tone made the strongest contributions to choice (Figure 3; note that tone-sequence lengths were distributed exponentially, with many consisting of just 2–3 tones, which may have limited our ability to detect effects for earlier tone positions).

**Figure 3.**
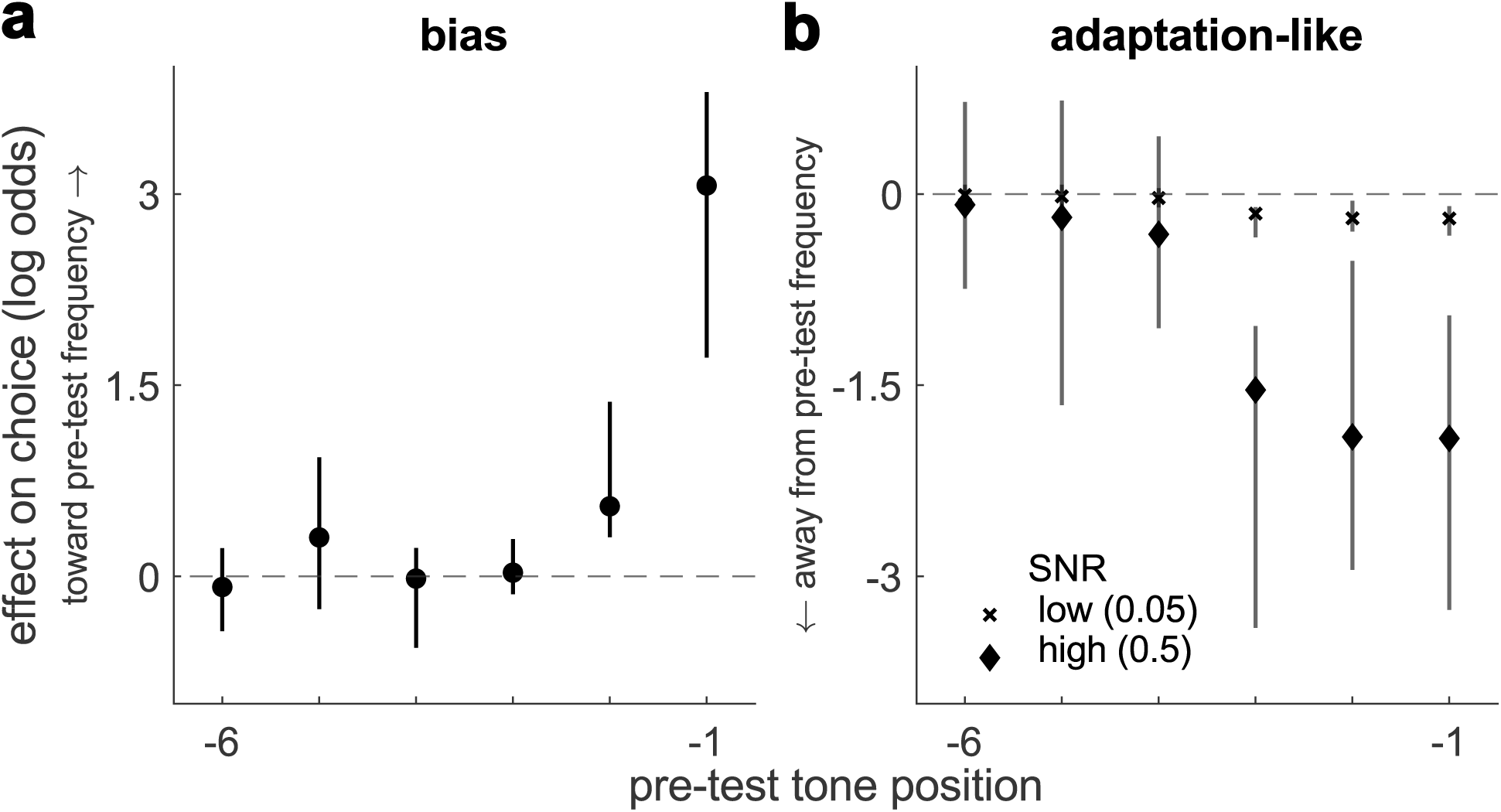
Logistic analysis of the bias (**a**) and adaptation-like (**b**) effects in the stimulus-based condition, as a function of pre-test tone position (−1 = the tone just prior to the test tone, etc.). Values along the ordinate are the medians of per-subject beta coefficients from the logistic model (points), and the error bars are bootstrapped 95% confidence intervals (computed using 10,000 bootstraps). These beta coefficients represent the effects of a pre-test tone presented at that position on choice, measured as log-odds ratios biasing choices toward (positive values) or away from (negative values) the frequency of the pre-test tone at the given position. In **b**, the two symbols (see legend) correspond to effects measured for different SNR values of the test tone. Note that the total effect of a given pre-test tone is found by summing the bias and adaptation-like effects, for a given test-tone SNR.

The negative SNR-dependent effect was unique to the stimulus-based condition: functionally analogous terms estimated in the rule-based condition, in which the effect of each cue was dependent on unsigned SNR, did not differ significantly from 0 (sign test, *p* > 0.05 for the SNR-dependent term for all three prior cues). A bias + adaptation model containing terms for the final two pre-test tone positions (Figure 2b, left; Table 1) was favored over both a bias-only model (mean ΔAIC = 3.11) and a model that did not include an effect of the tones (mean ΔAIC = 56.86). A Bayesian random-effects analysis confirmed the bias + adaption model as the most likely model (protected exceedance probability = 1). Though BIC favored the bias-only model (Table 1), it is known to be overly conservative in many cases [35, 36], and we further examined the necessity of the adaptation-like mechanism in subsequent analyses.

In summary, in the rule-based condition choices and RTs were both consistently biased in favor of the cue, and this effect diminished from low to high SNR in a manner that is consistent with normative principles. In contrast, in the stimulus-based condition choices and RTs exhibited a different pattern, which was characterized by a bias toward the cues at low test-tone SNR but a bias away from the cues at high test-tone SNR. Together, these results suggest that rule-based and stimulus-based cues may invoke different mechanisms.

### Rule- and stimulus-based biases were captured quantitatively by drift-diffusion model fits

The behavioral analyses detailed above indicate fundamental differences in how rule-based versus stimulus-based cues affect the speed and accuracy of auditory decisions. To better identify the computations responsible for these effects, we fit drift-diffusion models (DDMs) to the behavioral data from each task condition [38]. Unlike the regression-based approaches above, DDMs model mechanisms that can jointly account for choice and RT data in a unified framework. In the DDM, noisy evidence is accumulated to a pre-defined bound, which we modeled using five free parameters: 1) *drift rate,* which influenced the rate at which the test tone contributed to evidence accumulation; 2) *bound height,* which was the total evidence required to commit to a decision at the beginning of the trial; 3) *bound slope*, which determined the rate of a linear decline of the bound over the course of a trial (i.e., a “collapsing bound” that decreases in height over time), to account for urgency effects resulting from the time pressure to make the decision before the trial was aborted after 2–3 s; 4) *non-decision time,* which was the portion of total reaction time that was not determined by decision formation (e.g., sensorimotor processing); and 5) *lapse rate*, which quantified error rates for easily perceivable stimuli.

We added several cue-dependent terms to account for bias effects. For the rule-based condition, we used two additional free parameters per prior cue to account for biases in: 1) the rate of evidence accumulation, which we reasoned could account primarily for the strong bias-induced shifts in the psychometric function (Figure 4a, b); and 2) the starting point of the evidence-accumulation process, which we reasoned could account primarily for the strong vertical asymmetries in the RTs (Figure 4c, d). For the stimulus-based condition, we used three additional parameters: 1) a bias in the rate of evidence accumulation; 2) a bias in the starting point of the evidence-accumulation process; and 3) a shared time constant that describes the rapid temporal decay in the influence of each pre-test tone on the two biasing parameters (i.e., an exponentially weighted sum of biasing effects caused by each preceding tone; Figure 3a). Positive/negative values for the cue-dependent bias terms corresponded to more and faster choices toward/away from the dominant frequency of the pre-test tone sequence (i.e., of the exponentially weighted sum).

**Figure 4.**
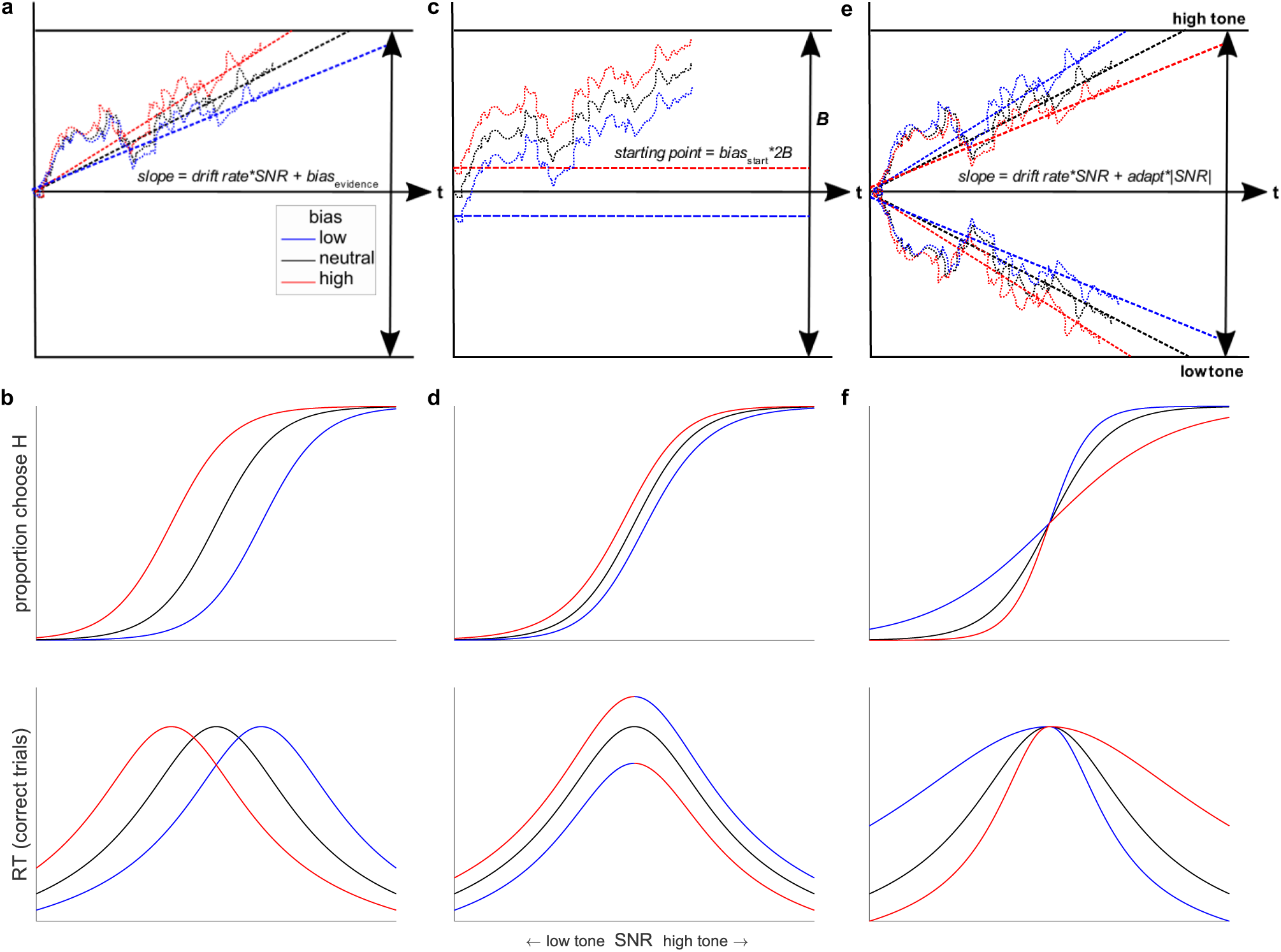
Simulated DDM bias terms (top) and their predicted, distinguishable effects on psychometric (choice; middle) and chronometric (RT; bottom) functions. **a** A bias implemented as an additive offset to the drift-rate term, which shifts the slope of the decision variable in the direction favored by the bias. **b** This “evidence-accumulation bias” causes lateral shifts in the psychometric and chronometric functions, corresponding to more and faster choices in the direction of the bias. **c** A bias implemented as a shift in the starting point of evidence accumulation toward the bound favored by the bias, implying that less evidence is needed to make a bias-congruent choice. **d** This “starting-point bias” causes smaller lateral shifts in the psychometric function than for the evidence-accumulation bias and vertical asymmetries in the chronometric function (see the discontinuity at SNR = 0), such that RTs are faster (slower) for prior congruent (incongruent) choices. **e** An adaptation-like bias that affects the slope of the decision variable in a manner that varies with the absolute value of the test-tone SNR. **f** For the stimulus-based condition, this term, which is absent at SNR = 0 and increases with increasing |SNR| of the test tone, shifts the slope of the decision variable away from the frequency of the tones heard prior to the test tone and thus causes more and faster choices to the frequency opposite the most-recently heard pre-test tones. **a, c, e** Dotted lines: example evolutions of the decision variable for a single trial with a given bias (see legend). Dashed lines: average slope of decision variable. *T* = time. In **c**, *B* = total bound height/2. **b, d, f** Simulated choice (top) and RT (bottom) data from an analytical solution to the DDM [37].

The starting-point and evidence-accumulation bias terms can produce effects that tend to diminish in magnitude with increasing unsigned SNR, as we saw in the rule-based condition. However, because these terms cannot produce the biases that go in the opposite direction at high SNRs, as we saw in the stimulus-based condition (see Figure 2b, d), we introduced a novel component to the model used for the stimulus-based condition, corresponding to SNR-dependent adaptation-like effects. Given the sharp decline in the influence of individual pre-test tones (Figure 3b) and previous work suggesting auditory neural adaptation effects have an exponential form [10, 39], we modeled the adaptation-like effect of the pre-test tones as an exponentially weighted sum of the effects of each tone (with a shared time constant but separate, multiplicative weights for high- and low-frequency tones), which was then multiplied by the SNR of the test tone. Negative weights imply that repeated pre-test tones caused more and faster choices away from the repeated tone frequency (Figure 4e, f). This mechanism, together with the bias terms, can produce the SNR-dependent effects that we found in the stimulus-based condition. Finally, we introduced an additional parameter to increase non-decision time as a function of the absolute value of the stimulus-based bias, which captured the tendency for faster RTs when the last two pre-test tones were the same frequency (Figure S1a; post-hoc test of interaction, *B =* 68.83, *t*(72.07) = 7.76, *p* < 0.0001).

The DDMs captured the patterns of bias and adaptation-like effects in the choice and RT data (Figure 5a, b, solid lines). In the rule-based condition, the “full” model with cue-dependent bias terms and a collapsing bound provided a better fit to the data than: 1) a model with no cue-dependent bias terms and a fixed bound (the “base” model); 2) a model with no cue-dependent bias terms and a collapsing bound (the “collapsing bound” model); and 3) a “full, fixed bound” model with cue-dependent bias terms but without a collapsing bound (Table 3; Figure S1b). However, there was also heterogeneity across subjects in the different forms of bias identified by the model fits. In particular, whereas the full model was the best-fitting model according to AIC and was estimated to be the most frequent model in the population (PEP = 0.79), an “evidence- only” model that was identical to the full model except for omitting the starting-point bias also received support (PEP = 0.20) and was considered the best model by BIC.

**Figure 5.**
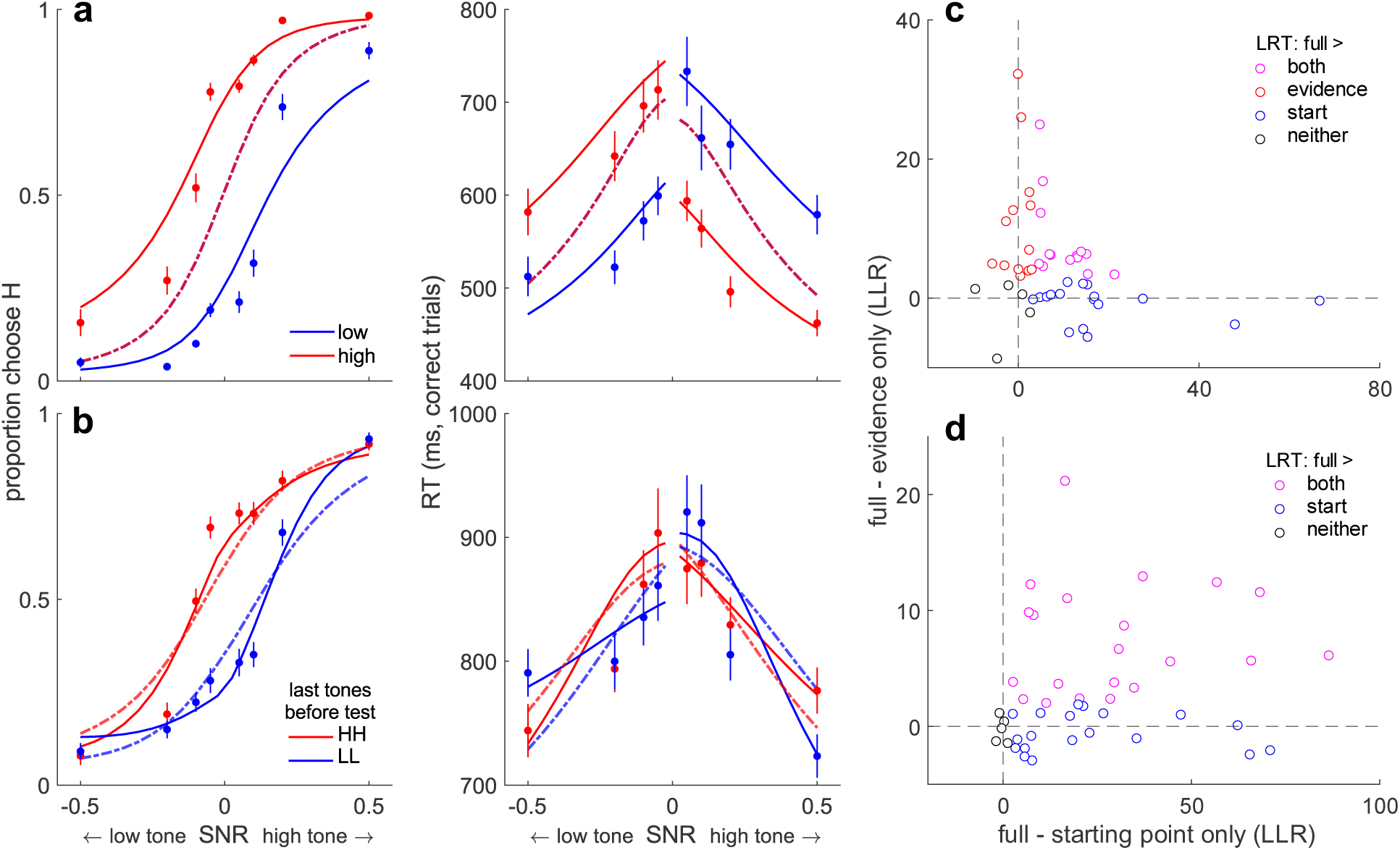
Comparing full and reduced model fits. **a** DDM fits to the rule-based condition for the base model, with no bias terms (dashed lines; note the overlapping lines), and the full model, with bias terms (solid lines). **b** DDM fits to the stimulus-based condition for the bias-only model (dashed lines) and full model (solid lines). **c** The difference in fit (log-likelihood ratio) for the rule- based full model vs. evidence-only model (ordinate) and full model vs. starting-point-only model (abscissa), plotted per subject (points). **d** same as **c**, for the stimulus-based condition. In **a** and **b**, points are averages over subjects’ data, and error bars are SEM. Fewer cue conditions are plotted than in Figure 2 to facilitate visual comparison between the full and reduced models. In **c** and **d**, data points are colored according to the model preferred from per-subject likelihood-ratio tests (LRT) between the full and reduced (evidence-only or starting-point-only) models (see legend). See Figure S2 for fits to the RT distributions for the full models.

**Table 3.**
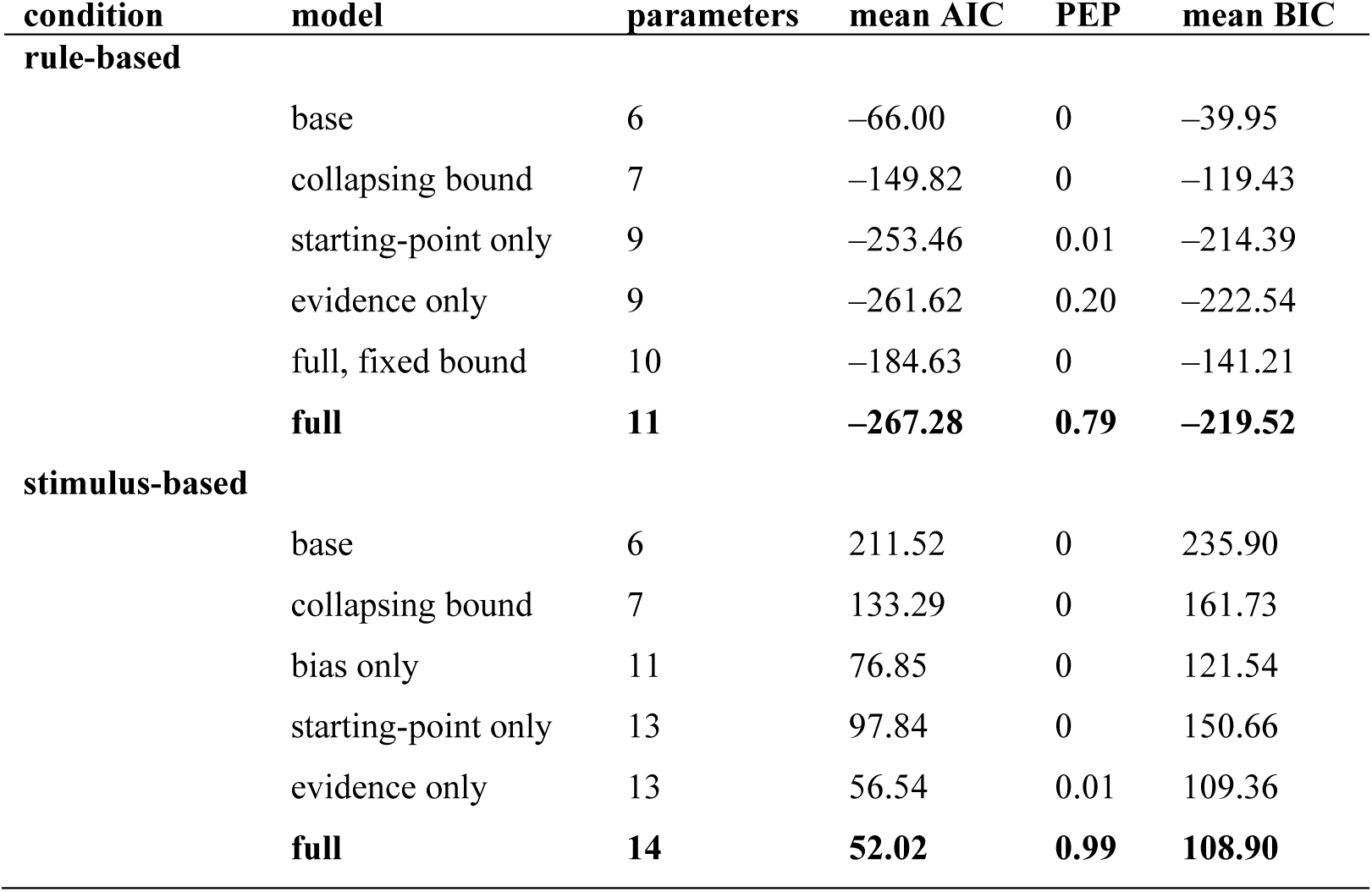
DDM model-fit comparisons. All models included evidence-acculumulation and starting-point bias terms to account for idiosyncratic biases, even when they did not include cue- dependent bias terms. See Methods for full implementation details. We selected best-fitting models (bold) on the basis of AIC and PEP.

| condition | model | parameters | mean AIC | PEP | mean BIC |
| --- | --- | --- | --- | --- | --- |
| **rule-based** |  |  |  |  |  |
|  | base | 6 | -66.00 | 0 | -39.95 |
|  | collapsing bound | 7 | -149.82 | 0 | -119.43 |
|  | starting-point only | 9 | -253.46 | 0.01 | -214.39 |
|  | evidence only | 9 | -261.62 | 0.20 | -222.54 |
|  | full, fixed bound | 10 | -184.63 | 0 | -141.21 |
|  | **full** | **11** | **-267.28** | **0.79** | **-219.52** |
| **stimulus-based** |  |  |  |  |  |
|  | base | 6 | 211.52 | 0 | 235.90 |
|  | collapsing bound | 7 | 133.29 | 0 | 161.73 |
|  | bias only | 11 | 76.85 | 0 | 121.54 |
|  | starting-point only | 13 | 97.84 | 0 | 150.66 |
|  | evidence only | 13 | 56.54 | 0.01 | 109.36 |
|  | **full** | **14** | **52.02** | **0.99** | **108.90** |

To better understand potential individual differences in the necessity of the bias terms, for each subject we computed likelihood-ratio tests comparing the full model to the evidence-only model and a “starting-point-only” model (i.e., omitting the evidence-accumulation bias). This analysis confirmed considerable heterogeneity across subjects (Figure 5c): 29% were better fit by the full model over both reduced models, with most of the remaining subjects split between being better fit by the full model over the starting-point-only model alone (35%) and the full model over the evidence-only model alone (27%; *p*s < 0.05). This pattern suggests that different subjects relied on starting-point biases and evidence-accumulation biases to differing degrees (with slightly more favoring an evidence-accumulation bias), an idea we examined in more detail below (see Figure 6). To encompass this variability, we used the full model in subsequent analyses.

**Figure 6.**
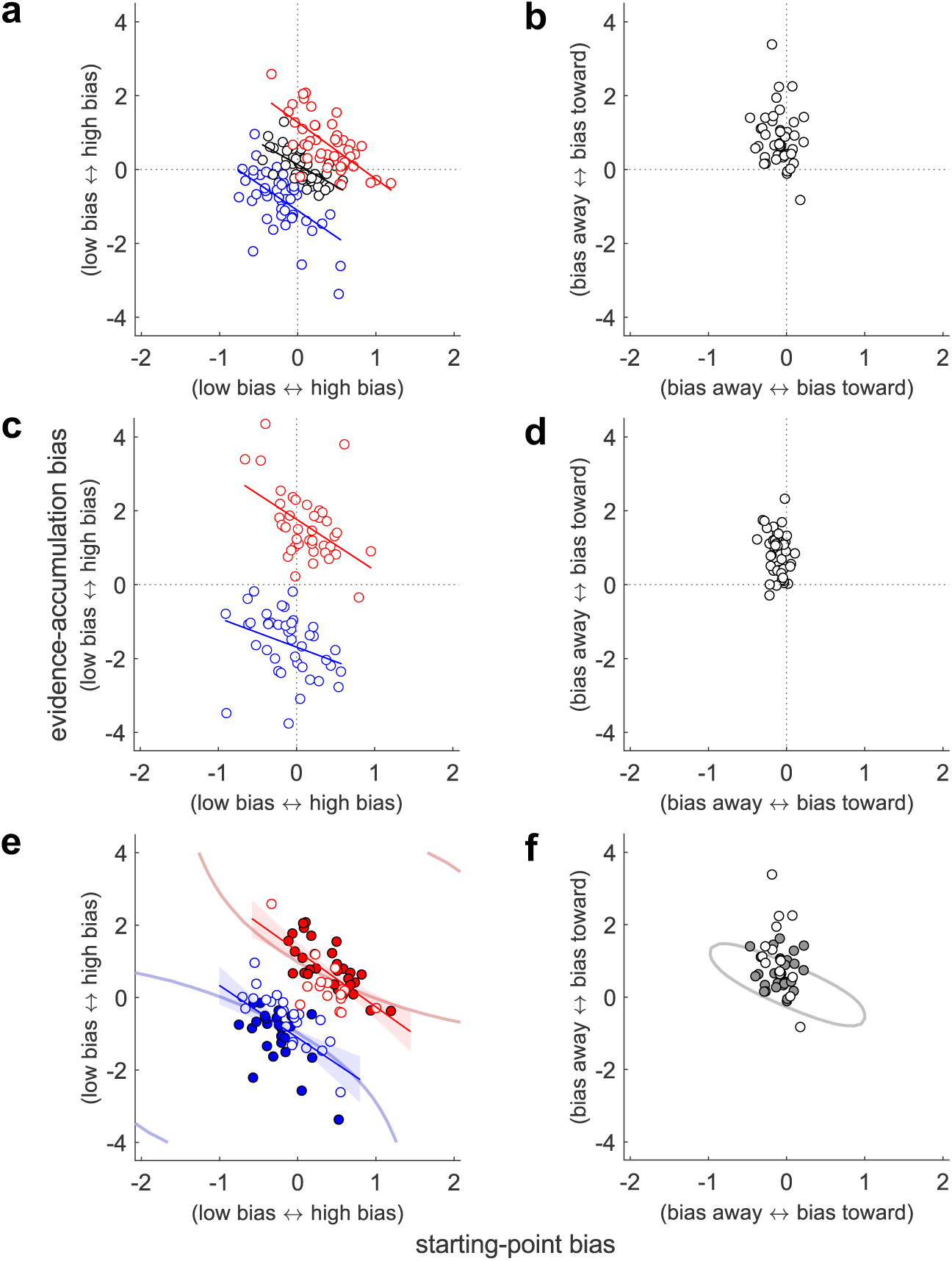
Different computational signatures of rule- and stimulus-based biases. Each panel shows best-fitting evidence-accumulation (ordinate) and starting-point biases (abscissa), plotted per subject (points). **a** Data from the rule-based condition, for each of the three prior cues (blue: low, black: neutral; red: high). Lines are linear fits (*p* < 0.05). **b** Data from in the stimulus-based condition (no linear relationship). **c** Data from the rule-based biases of the mixed condition, plotted as in **a**. **d** Data from the stimulus-based biases of the mixed condition, plotted as in **b** (no linear relationship). **c** and **d** show results for mixed-condition 1, which included block-level prior cues; results were similar in mixed-condition 2, which had trial-wise priors. **e** Data from **a** (the low- and high-prior conditions) plotted with respect to optimal performance contours from the rule-based condition. The average 97% contours (curved lines) demarcate the area of the parameter space in which choice accuracy is at least 97% of the maximum possible accuracy. Accuracy was predicted by the DDM applied to the actual task sequences. Closed/open circles indicate that the data fell within/outside of the subject-specific 97% contour. Lines are linear fits as in **a**, and shaded regions are 95% confidence intervals. **f** Same as **e**, but for the stimulus-based condition. The 97% contour is for the context of two low tones before the test tone (similar results were obtained for two high tones but not shown).

In the stimulus-based condition, the “full” model with a collapsing bound and both bias and SNR- dependent terms provided a better fit to the data than a base model without biases, a collapsing-bound model without biases, and a collapsing-bound model with only bias terms but no SNR- dependent (i.e., adaptation-like) terms (Table 3). Not only did the “bias-only” model fail to capture the crossover interaction from low to high SNR (compare solid [full model] versus dashed [bias- only model] lines in Figure 5b), but the SNR-dependent terms in the best-fitting full models were consistently negative, confirming an adaptation-like effect (one-sample *t*-test; both *p*s_corrected_ < 0.0001).

Additionally, the full model was favored over both an evidence-only and a starting-point-only model (Table 3), suggesting the necessity of both mechanisms. To confirm this result and assess heterogeneity across subjects, we performed an equivalent likelihood-ratio-test analysis to the rule- based condition (Figure 5d). Whereas this analysis supported the full model overall (47% of subjects were fit better by the full model over both reduced models), 42% of subjects were better fit by the full model over the starting-point-only model but not the evidence-only model (*p*s < 0.05). This result suggests that the evidence-accumulation bias was necessary to account for performance across subjects, but an additional starting-point bias was necessary only for about half of the individual subjects. Given that the full model appears to be most frequent in the population (PEP = 0.99) and subsumes the evidence-only model, we primarily used the full model in subsequent analyses (and note when otherwise).

### Rule- and stimulus-based biases exhibited distinct computational signatures

The regression analyses and DDM fits together demonstrate that the stimulus-based effects, unlike the rule-based effects, require an adaptation-like mechanism. However, these analyses leave open the question of whether the remaining biases that were present in both conditions (i.e., those that tended to produce more and faster choices in the direction of the rule-based cues at all SNRs and of the stimulus-based cues at low SNRs) rely on the same or distinct mechanisms. To address this question, we examined computational signatures of the evidence-accumulation and starting-point biases derived from the DDM fits.

For the rule-based condition, subjects tended to use both starting-point and evidence-accumulation biases in the direction of the higher-probability alternative (Table 4, light green rows; one sample *t*-test, all *p*s_corrected_ < 0.0001). Across subjects, these two kinds of computational biases were negatively correlated with one another: subjects who tended to use stronger starting-point biases had weaker evidence-accumulation biases and vice versa. This negative relationship was evident within all three prior cue conditions (Figure 6a; low: Spearman’s *ρ* = −0.49, *p*_corrected_ = 0.0003; neutral: *ρ* = −0.72, *p*_corrected_ < 0.0001; high: *ρ* = −0.62, *p*_corrected_ < 0.0001) and mirrors the heterogeneity in goodness-of-fit across subjects (Figure 5c).

**Table 4.**
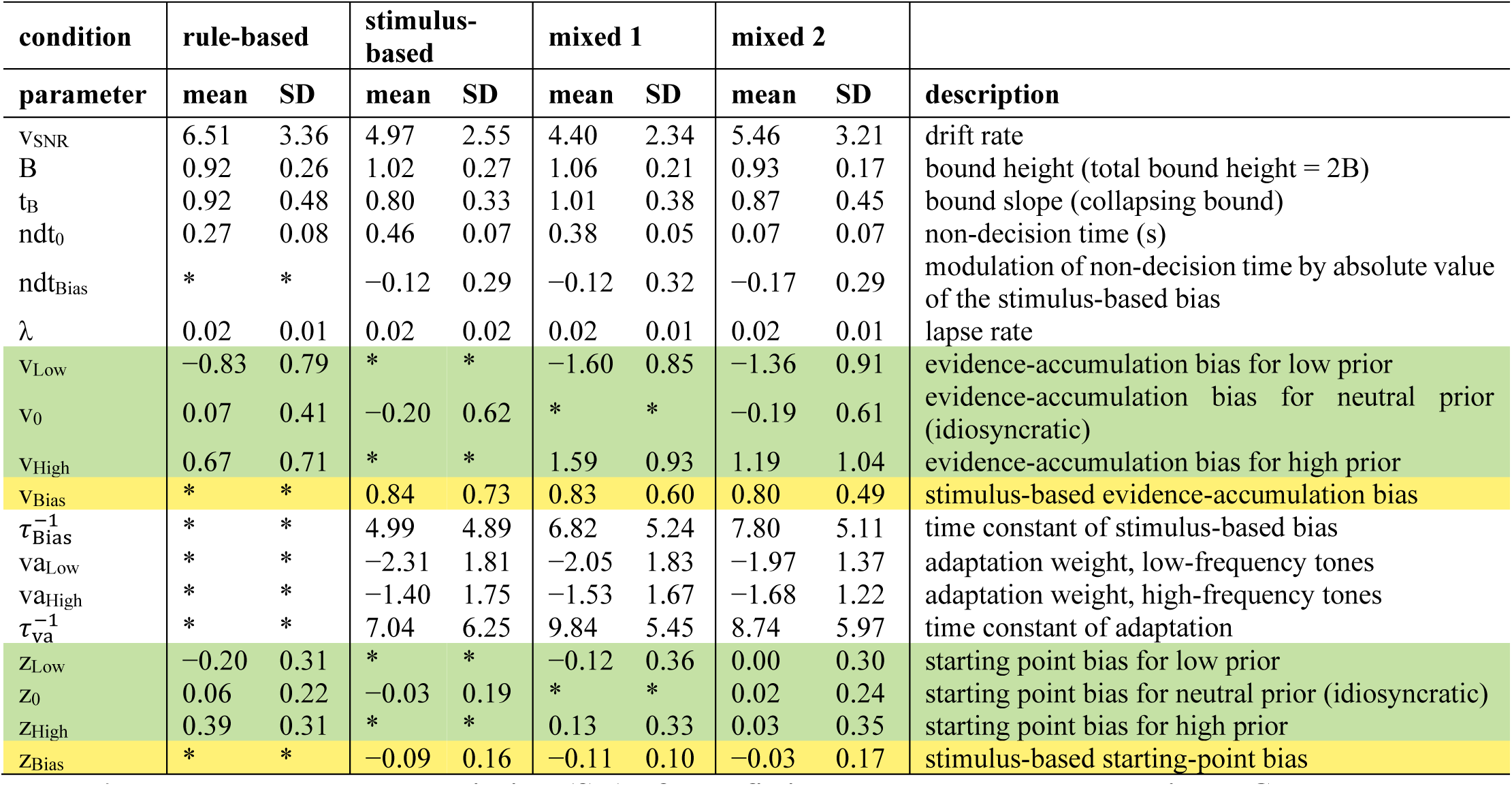
Mean and standard deviation (SD) of best-fitting paramaters across subjects. . Green rows: rule-based bias parameters; yellow rows: stimulus-based bias parameters; * parameter not present in that condition. See Methods for a fuller description of the parameters.

For the stimulus-based condition, subjects also tended to use a combination of starting-point and evidence-accumulation biases but with a different pattern than for the rule-based condition (Table 4, light yellow rows). Specifically, the evidence-accumulation bias tended to be positive (i.e., toward the frequency of the most recent tones; one-sample *t-* test, *p* < 0.0001). In contrast, the starting-point bias tended to be in the opposite direction, away from the bound for the frequency of the most recent tones (*p* = 0.0007). Unlike for the rule-based DDM fits, we could not identify any across-subject correlation between best-fitting values of evidence-accumulation and starting-point biases (Figure 6b; Spearman’s *ρ* = −0.20, *p* = 0.19). In the two mixed-condition conditions, we confirmed that this differential pattern of correlations was also present and thus was not simply an idiosyncratic quirk of either the rule- or stimulus-based condition tested individually (rule-based: Figure 6c, all *p*s_corrected_ < 0.005; stimulus-based: Figure 6d, all *p*s > 0.05).

The negative relationship between evidence-accumulation and starting-point biases in the rule-based conditions is similar to that found for monkeys performing a reward-biased visual-decision task [40, 41]. In that study, the reward-driven relationship was interpreted in terms of coupled, goal-directed (top-down) adjustments to the decision process that improved performance. Specifically, monkeys made adjustments in evidence-accumulation and starting-point biases that were sensitive to session- and monkey-specific variations in perceptual sensitivity and contextual factors. These adjustments maintained near-optimal or “good enough” (satisficing) performance, possibly by using a learning process to rapidly adjust the biases until reaching a performance plateau. Critically, the contours of this performance plateau showed a “tilt” in the space of evidence-accumulation and starting-point biases, such that negatively correlated values of these biases traced out iso-performance contours. We reasoned that a similar sensitivity to performance- satisficing combinations of biases could plausibly underlie the correlation between the evidence-accumulation and starting-point biases in our rule-based condition.

To assess this possibility, we used the DDM to predict, for each subject, the average choice accuracy that could be achieved across a range of evidence-accumulation and starting-point bias parameters, given the subject’s other fit parameters and the task context. For the rule-based condition, task context was determined by the prior cue. For the stimulus-based condition, task context was defined with respect to two high or two low pre-test tones, which had the strongest effects on behavior in that condition. To facilitate comparisons, we normalized predicted accuracy for each subject and condition to the maximum value achieved via simulations, using a wide range of evidence-accumulation and starting-point bias parameter values.

For the rule-based condition, the subjects’ combinations of evidence-accumulation and starting-point biases tended to yield nearly optimal performance (low: median proportion maximum performance = 0.97, range = [0.76–1.00]; neutral: 1.00 [0.97–1.00]; high: 0.98 [0.86–1.00]). Furthermore, the location and tilt of the across-subjects distribution of biases appeared qualitatively to respect the tilt of the plateau of the performance function (Figure 6e), suggesting that sensitivity to this plateau was a potential cause of the correlation between the biases. For the stimulus-based condition, the subjects’ biases also yielded nearly optimal performance (LL: median proportion max performance = 0.99, range = [0.62–1.00]; HH: 0.99 [0.80–1.00]), reflecting the fact that some degree of positive bias was needed to offset performance decrements induced by the adaptation-like effect. In contrast, the subjects’ biases for the stimulus-based condition did not follow the tilt of the plateau of the performance function (Figure 6f).

To confirm this pattern of results, we used a mixed-effects model to compare the slopes of the linear fit that predicted the starting-point bias from the evidence-accumulation bias between the different contexts. We could not identify any difference in the slopes of the high versus low biases within the rule-based condition (*B =* −0.03, *t*(126.00) = −0.14, *p* > 0.05). In contrast, there was a significant difference in the slope of the biases between the rule-based and stimulus-based conditions (*B =* 0.45, *t*(126.00) = 2.80, *p =* 0.006). These findings are consistent with the idea that, at low test-tone SNR, the subjects’ choices in the stimulus-based condition were produced by a different underlying mechanism than the coupled (tilted) changes that mediated choices in the rule-based condition.

Further supporting this idea of different mechanisms, we could not find any evidence that individual subjects made consistent use of starting-point and/or evidence-accumulation biases across the rule-based and stimulus-based conditions. Specifically, there was no correlation between best-fitting values of each bias term computed per subject when comparing across the two conditions (*p* > 0.05 for both starting-point and evidence-accumulation biases). When rule-based and stimulus-based biases were used at the same time in the mixed conditions, we also could not identify a correlation between the bias terms (*ps* > 0.05 for both starting-point and evidence-accumulation biases in both mixed conditions; note that parameter ranges were similar for the single and mixed blocks, suggesting that the subjects used the biases in a roughly comparable, although not identical, way in the two block types; Table 4).

Together, these results suggest that rule-based and stimulus-based biases rely on separable computational processes. One possible caveat to the conclusions of the preceding analyses arises from the possibility that the start-point bias may be unnecessary to account for the stimulus-based effects, hence it did not correlate with the evidence-accumulation bias because it only picked up noise. However, this outcome would still be strong evidence for differences between the rule- and stimulus-based biases, given that both biases were necessary to account for the range of individual differences in the rule-based condition. Furthermore, the evidence-accumulation bias from the stimulus-based evidence-only model was uncorrelated with the rule-based evidence-accumulation bias (*p* > 0.05), again supporting the idea that the computational basis for the behavioral biases differed for the two task conditions.

### Rule- and stimulus-based biases exhibited distinct physiological signatures

Because non-luminance-mediated changes in pupil size can be associated with particular forms of behavioral decision biases [9,21,29–31], we tested whether these pupil changes could be used to further distinguish the different effects of rule-based and stimulus-based cues on auditory decision-making that we identified above. In particular, if pupil-linked arousal facilitates overcoming biases, as has been proposed previously [9, 30], then we would expect larger evoked pupil responses when comparing incongruent to congruent trials: responding correctly on incongruent trials requires overcoming bias. Furthermore, if the rule- and stimulus-based biases depend on overlapping neurophysiological mechanisms, then we would expect similar profiles of pupil responses for both conditions. In contrast, if pupil-linked arousal for our task is more sensitive to top-down than to bottom-up influences, as has also been reported previously [32], then we would expect more strongly modulated pupil responses in the rule-based than in the stimulus-based condition.

In the rule-based condition, the choice-aligned evoked pupil response was modulated strongly by congruence (Figure 7a, b). Specifically, this pupil response was larger for correct choices that were incongruent with the prior relative to those that were congruent with the prior. This effect emerged shortly after choice (because the effects were similar but weaker when aligned to stimulus onset, we focus on choice-aligned effects). We found a similar effect on incorrect trials (Figure 7d, e), which implies that these pupil modulations were more closely related to the congruence between the prior and the choice than to the congruence between the prior and the stimulus. We also found that pupil size was modulated on both congruent and incongruent trials relative to neutral trials, with larger dilations on neutral trials starting before stimulus onset. These dilations likely reflected higher uncertainty on those trials (i.e., a less-informative prior; Figure 7a, b). In the mixed conditions, similar rule-based modulations were also evident and thus were robust to the presence or absence of stimulus-based biases (mixed-effects models comparing average pupil responses 220–720 ms post-choice for pairs of prior cues, all *p*s_corrected_ < 0.03, except for the neutral– incongruent contrast in mixed block 2, *p* > 0.05).

**Figure 7.**
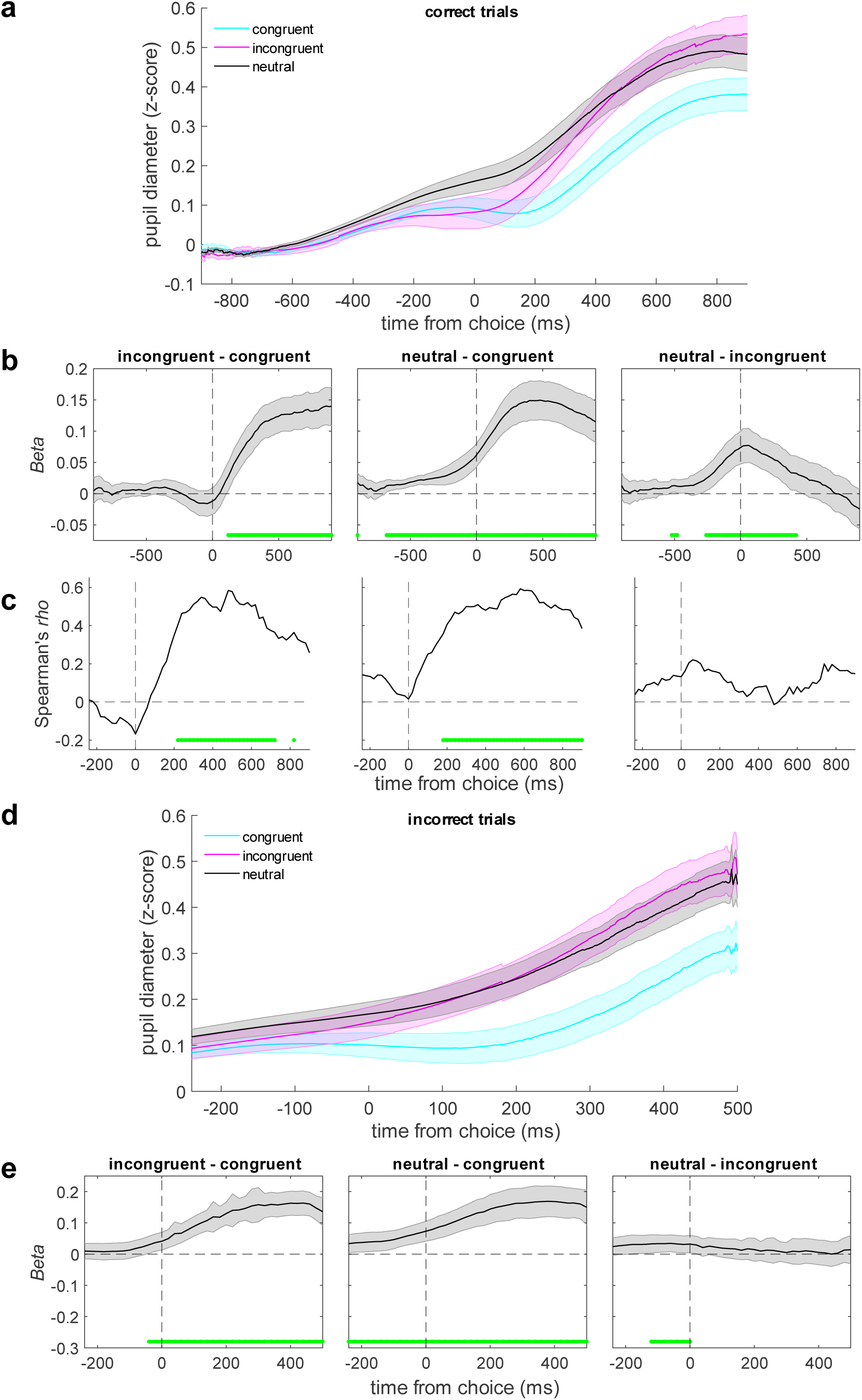
Rule-based cues reliably modulated pupil-linked arousal. **a** Pupil diameter on correct trials, aligned to choice and plotted as a function of the congruence between the choice and the prior. Lines and error bars are means and SEM across subjects. **b** Regression coefficients (beta values) from a linear mixed-effects model as a function of different pupil contrasts (see labels above each panel). Error bars are 95% confidence intervals of the parameter estimates. **c** Across- subject Spearman correlation between: 1) the degree to which each subject was biased by the prior cues, estimated from a logistic regression model of choice behavior; and 2) each subject’s pupil contrast from a linear regression model. **d, e** Same as **a, b** for incorrect trials. The time window in **d** and **e** was chosen to focus on the period in which the incongruent – congruent effect was most prominent in the correct trials (and note that incorrect trials ended earlier than correct trials; see Methods). Values in **b**, **c,** and **e** were computed in 20 ms-wide time bins. Green lines indicate time bins for which *p*_corrected_ < 0.05, FDR-corrected across time and contrast.

The magnitude of these rule-based pupil modulations was correlated with the magnitude of behavioral biases across subjects (Figure 7c). Specifically, we measured the Spearman correlation coefficient between the magnitude of choice bias derived from the logistic fits to each subject’s choice data (high-prior-cue bias minus low-prior-cue bias) and pupil contrast (β value). We found positive correlations (∼200–800 ms post-choice) for both the congruent–incongruent and neutral– congruent contrasts. In other words, a greater reliance on the prior cues corresponded to more differentiated arousal responses between trials when the cue was or was not congruent with the choice. Our results complement earlier findings that pupil responses can reflect the magnitude of (inappropriate) choice biases, with larger pupil responses typically corresponding to a reduction in bias [9,21,29,30]. These results suggest that our subjects, particularly those who were highly sensitive to the prior cues, needed to mobilize additional resources to overcome the prior or make decisions in the absence of an informative prior.

In contrast, evoked changes in pupil size in the stimulus-based condition displayed a very different pattern of results. Because the subjects’ behavior in this condition depended on congruence and SNR, we analyzed pupil size as a function of the interaction between: 1) the congruence of choice with the last two pre-test tones and 2) SNR. We analyzed only those trials in which the pre-test tones were the same frequency and the test-tone SNR was lowest or highest, to increase our chances of finding effects. However, unlike for the rule-based condition, we did not identify any effects that survived correction for multiple comparisons (Figure 8). Further, we found a significant congruence×condition interaction (mixed-effects model comparing average pupil responses 0–1000 ms post-choice, *B =* 0.09, *t*(38.36) = 5.65, *p* < 0.0001), which confirmed that the congruence-related arousal response was stronger in the rule-based condition than in the stimulus-based condition. Although we could not identify any sign of stimulus-based effects in the post-choice period, we found a congruence×SNR interaction at an uncorrected *p* < 0.05 from 740– 220 ms before choice. We also reanalyzed the rule-based condition to look for a congruence×SNR interaction in that time window and could not identify any effect (all *p*s > 0.05).

**Figure 8.**
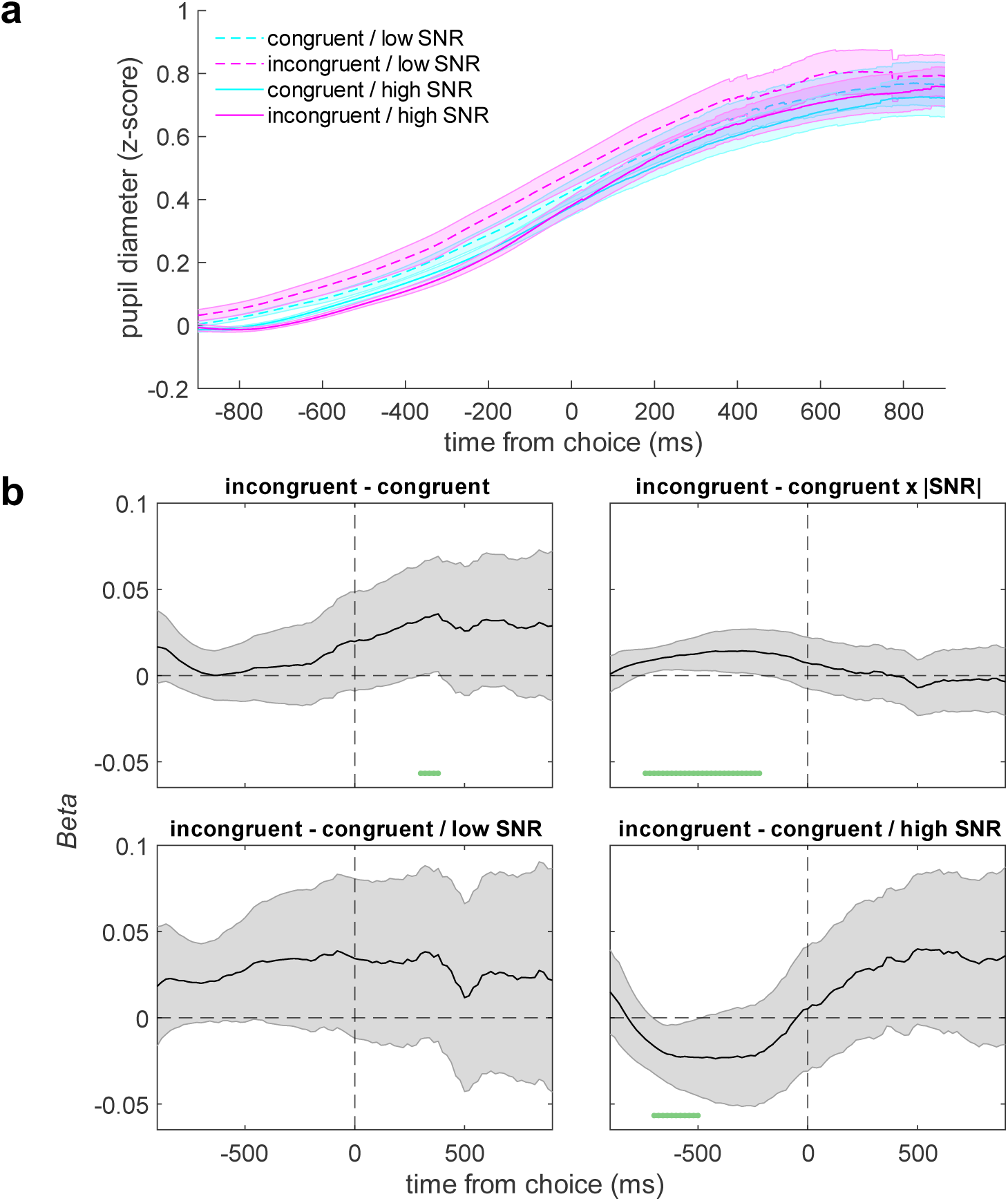
Stimulus-based cues weakly modulated pupil-linked arousal. **a** Pupil diameter on correct trials, aligned to choice and plotted as a function of the congruence of the choice and the most-recently heard pre-test tones, separated by SNR. We only used pre-test sequences ending in two of the same tone (i.e., LL, HH) and at the lowest or highest unsigned test-tone SNR for this analysis. Lines and error bars are means and SEM across all trials and subjects. **b** Regression coefficients (beta values) from a linear mixed-effects model for different pupil contrasts. Error bars are 95% confidence intervals of the parameter estimates. Values were computed in 20 ms- wide time bins. Dark green lines indicate time bins for which *p*_uncorrected_ < 0.05.

In summary, pupil-linked arousal strongly differentiated between the rule- and stimulus-based biases in the locus, strength, and timing of the effect. In the rule-based condition, pupil size following choice was robustly indicative of the congruence of the choice and the prior cue in a manner that reflected individual behavioral biases. In the stimulus-based condition, pupil size preceding choice only weakly reflected the SNR-dependent choice-congruence effects.

## Discussion

This work was motivated by the realization that, despite an abundance of studies on the importance of expectations in auditory processing, we lack a basic understanding of their effects on perception. We focused on how human auditory perceptual decisions are influenced by two different sources of expectations, one from instructed rules that indicated the probability of the upcoming test tone and the other from stimulus regularities akin to those used in oddball tasks. Rule-based cues consistently biased behavioral choices and RTs in favor of the more probable alternative. In contrast, stimulus-based cues elicited different biases that depended on the test-tone SNR. We leveraged model fits and measures of pupil-linked arousal to show that the rule-based and stimulus-based behavioral biases have distinct computational and physiological signatures.

Our results extend previous findings in three fundamental ways. First, we were able to decompose the effects of stimulus regularities on auditory perception into an adaptation-like effect on stimulus sensitivity and a decision bias, whereas prior work in auditory perception tended to focus on only one or the other of these effects. Second, because of this decomposition, we could directly compare decision biases produced by rule-based versus stimulus-based cues. We found that rule-based and stimulus-based biases have distinct computational signatures. In particular, the effects on the decision variable (i.e., evidence-accumulation bias in the DDM) and the decision rule (i.e., starting-point bias in the DDM) are coupled for rule-based biases but not for stimulus-based biases. Third, we showed that rule-based and stimulus-based biases also have distinct physiological signatures: pupil-linked arousal is modulated by rule-based biases but not by stimulus-based biases. As detailed below, we interpret these results in terms of two potentially distinct mechanisms for incorporating prior expectations into auditory perceptual decisions: one using top-down processing for rule-based cues and the second using bottom-up processing for stimulus- based cues.

The rule-based cues in our task were visual, not auditory, and thus were, at some level, processed separately from the incoming auditory stream and then integrated into the auditory decision. Several of our findings imply that this process involved cognitively driven, top-down mechanisms but not lower-level interactions between the visual and auditory stimuli [42, 43]. We modeled the rule-based decision biases in terms of computational components of a normative, DDM-based decision process. These kinds of computationally defined biases, which have been studied extensively for visual decision-making [4–7,44,45] but only sparsely for auditory decision-making [1, 9], have been ascribed to top-down influences [8]. Our results further support this idea by showing that auditory decision biases can occur flexibly, depending on the relationship between the cue and the test stimulus. Moreover, these biases involve computational components that are coordinated with each other, which we previously showed is consistent with a cognitive-based (i.e., not purely stimulus-driven) learning process that can allow individual decision-makers to achieve nearly optimal performance [40].

Likewise, the rule-based biases that we measured were accompanied by modulations of pupil size that, like for related findings, appear to be cognitively driven [28]. Specifically, we found transient pupil increases in response to choices that were incongruent with the prior cue on that trial. These pupil modulations were similar to those that have been reported for other tasks involving decision biases [9,21,29–31], violations of learned expectations [32], and certain task-relevant but not task- irrelevant stimulus changes [46], all of which were interpreted in terms of arousal-linked cognitive influences on perception and/or decision processing. Our work complements and extends these findings by showing that the rule-based effects on task behavior and pupil size can be adjusted, based on the current version of the cue and its relationship (congruence) with the test tone, further supporting the idea that they are driven by the kinds of flexible information processing associated with top-down control.

Together, these results are consistent with the idea that arousal-linked processes actively monitor and adjust decision-making using relevant predictive cues. These adjustments likely arise from multiple neuromodulatory systems, principally the locus-coeruleus (LC) norepinephrine system and basal forebrain cholinergic system [28,47,48], that can affect either bottom-up or top-down information processing in the brain under different conditions. For example, relatively slow fluctuations in baseline or “tonic” arousal levels have been linked to modulation of sensory neurons [49–55] and changes in perceptual sensitivity in animal models and human subjects [49,56,57], suggesting bottom-up effects. In contrast, relatively fast, event-driven “phasic” changes in arousal have inconsistent relationships with perceptual sensitivity in human subjects [9, 29]. Rather, these phasic changes (like what we measured) are more strongly associated with behaviorally relevant decision- and response-related processes in both humans and animal models [30,58–60], particularly those requiring cognitive or behavioral shifts [61], which is suggestive of top-down effects.

Our results support this interpretation: the pupil response was modulated by the congruence between the cue and the subject’s choice, not the cue and the tone identity, consistent with arousal-linked adjustment of top-down decision biases rather than bottom-up sensory processing. This interpretation accords with a previous visual decision-making study showing that pupil-linked arousal modulates decision biases in frontoparietal regions encoding choice, without any effect on visual cortex [29]. Further work is needed to identify whether similar neural dissociations exist in auditory decision-making.

In contrast, the stimulus-based cues appeared to engage a different set of mechanisms that were more consistent with contributions from bottom-up processing. In particular, the stimulus-based cues elicited behavioral biases that included both “attractive” effects (i.e., biases towards previous stimuli) and “repulsive” effects (i.e., biases away from previous stimuli). Both of these effects have been identified in previous studies of auditory perception [3,15,17,18,62–65] and auditory short-term memory [66]. However, most of these prior studies reported consistently attractive or repulsive effects (cf. [65] for opposing effects of prior stimuli and prior perception) but not both, leaving us without a principled explanation for the conditions under which one or the other effect should occur.

To our knowledge, we are the first to report that the direction of bias depends strongly on SNR, with attractive decision biases predominating for low-SNR test stimuli and repulsive, adaptation- like mechanisms that reduce sensitivity to repeated (congruent) stimuli and increase sensitivity to deviant (incongruent) stimuli predominating for high-SNR test stimuli. One possible explanation for this SNR-dependence is that the recovery from adaptation, which is responsible for the increased sensitivity to incongruent test tones, is much weaker at low SNR, due to the more broadband nature of the stimulus. To capture the repulsive effects, which could not be accounted for with standard bias terms, we introduced a novel SNR-dependent adaptation component to the DDM. Though the attractive stimulus-based biases were well-characterized by DDM components similar to those for the rule-based effects, they were unrelated to rule-based biases and did not show any coordination within subjects. Moreover, the stimulus-based effects had little relationship to pupil size, unlike the rule-based effects. Thus, these effects did not appear to engage the top- down, arousal-linked cognitive mechanisms used to generate rule-based biases but instead were induced by the tone sequence itself.

We propose that the stimulus-based effects were likely driven, in part, by stimulus-specific sensory adaptation. Stimulus-specific adaptation affects neural responses throughout the auditory system and generally involves reduced responses to repeated presentation of the same stimuli and potentiated responses to novel (deviant) stimuli [10,26,27,34]. There is debate over whether these phenomena reflect (top-down) predictions or whether they are indicative of traditionally construed bottom-up sensory adaptation, with evidence suggesting that both processes may contribute [2,10– 14,34,67]. Both interpretations are consistent with our findings, in that the attractive effects could reflect predictions whereas the repulsive effects are more consistent with an adaptation-like process.

A possible reason for the coexistence of both stimulus-based attractive and repulsive effects is to allow the brain to navigate the tradeoff between stable integration and change detection as a function of the quality of sensory evidence [68, 69]. When SNR is low, the evidence is unreliable and thus prior expectations are useful. In contrast, when SNR is high and prior expectations are not as useful, other aspects of stimulus processing can be prioritized, such as increasing sensitivity to deviations from stimulus regularities. Past normative models of evidence accumulation in changing environments have accounted for human subjects’ sensitivity to change via a volatility-dependent leak on the prior accumulated evidence [68]. Future work should investigate whether adaptation-induced changes in the encoding of sensory evidence could also account for subject behavior [16,69–71], and whether environmental volatility might alter the balance between attractive and repulsive effects in auditory decision-making.

Though we believe the repulsive effects are consistent with stimulus-specific adaptation, it is possible that other cognitive or neural mechanisms could explain the patterns in the data. For example, another phenomenon that can produce faciliatory effects for deviant stimuli is “pop-out” [72, 73], in which a unique target stimulus is rapidly identified when presented amidst a group of identical stimuli that differ from the target on some key feature(s) (e.g., in the stimulus sequence HHL, where L is the test tone). However, whereas pop-out predicts a purely facilitatory effect of incongruent tones, adaptation predicts both a facilitatory effect of incongruent tones and an inhibitory effect of congruent tones, which is what we found (e.g., compare the blue fit lines at SNR = −0.5 and the red fit lines at SNR = 0.5 in Figure 5b, right, which show a repulsive effect in the data and the full model but not the bias-only model). Furthermore pop-out seems unlikely to explain the similar effects found in non-repeating pre-test tone trials (i.e., HL, LH), in which there is presumably nothing to pop-out because there was no repeated tone.

More generally, our models captured key qualitative patterns in the data but at times deviated quantitatively from the behavior of human subjects. Though we conducted careful model comparisons, it is always possible that alternative or additional mechanisms other than those considered here could better account for the data. Given the complexity of our models and the limits of our data, we erred on the side of parsimony in not considering further model components. Future research should build upon this work to delineate the computational bases of rule- and stimulus-based behavioral biases more precisely.

In sum, the results of this study established new methods for understanding how stimulus- and rule-based information shape auditory decisions, jointly within a DDM framework. Future work should leverage this framework to identify the neural substrates of these effects and should further establish the stimulus dimensions that govern the balance between attractive and repulsive effects, which has implications for understanding how the brain balances reliance on prior information with sensitivity to change.

## Materials and methods

Fifty human subjects participated in the study (19 male, 25 female, 6 N/A; median age: 25 yrs, range 18–60). Informed consent was obtained in accordance with the University of Pennsylvania IRB.

### Behavioral task

Each subject was seated and faced an LCD computer screen (iMac), which sat on a table in a darkened room. The subject listened to auditory stimuli via earphones (Etymotic MC5), while a chin rest stabilized their head for eye tracking. The subject reported whether a “test” tone burst (250 or 2000 Hz; 300-ms duration; 10-ms cos^2^ ramp) was “low” frequency or “high” frequency by pressing a button on a gamepad with their left or right index finger, respectively. This test tone was embedded in a background of broadband noise (50–20000 Hz), and we titrated task difficulty by varying the sound level of the test tone (low frequency: 53, 57, 62, and 69 dB SPL; high frequency: 52, 57, 62, and 69 dB SPL) relative to the noise (75 dB SPL), producing signal-to-noise ratios (SNRs) of −23 to −6 dB. Because the DDM expects SNR to be a positive quantity that increases with numerical magnitude, we mapped these SNR values onto an (unsigned) scale between 0.05 – 0.5. We also explored alternative SNR mappings, such as linearly rescaling dB SNR between 0.05 and 1.0, and the alternatives produced nearly equivalent results when fitting the psychometric function. Auditory stimuli were generated in the Matlab (R2016a) programming environment.

Most subjects completed 4–5 sessions of the task. Each session began with a set of 60–120 training trials. We then tested a different set of expectation-generating cues per session: “rule-based” cues, “stimulus-based” cues, or both (“mixed” conditions; see Figure 1a). In a rule-based-cue session (576 trials), we presented a visual cue at the beginning of each trial: a triangle pointing to the left indicated 5:1 odds of a low-frequency test tone, a triangle pointing to the right indicated 5:1 odds of a high-frequency test tone, and a square indicated even (neutral) odds. The same cue was presented for a block of 48 trials and was varied randomly between blocks.

In a stimulus-based-cue session (480 trials), we presented a tone-burst sequence (250 or 2000 Hz; 300-ms duration; 10-ms cos^2^ ramp; 100-ms inter-burst interval) that preceded the test tone (i.e., the “pre-test” sequence). This pre-test sequence was always presented at the highest SNR. The length of the pre-test sequence varied between 2–14 tones (approximately exponentially distributed). These sequences were pseudorandomly generated, but we ensured that, in each session, there were approximately the same number of high- and low-frequency tones.

We also had mixed sessions in which subjects had both rule-based and stimulus-based cues on each trial. In mixed-condition 1, which was divided into two sessions, we used the same rule-based cue (high or low prior) for the entire session (480/trials session) and varied the length of the pre-test sequence. In mixed-condition 2 (576 trials), we randomly varied the rule-based cue and fixed the length of the pre-test sequence at five tone bursts.

Sessions were completed in the following order: rule-based, stimulus-based, mixed-condition 1 (using either the low- or high-prior cue, randomly selected), mixed-condition 1 (using the alternative prior cue), and mixed-condition 2. Because of attrition and other factors, not all subjects completed all conditions: rule-based, *N*=49 subjects completed the condition; stimulus-based, *N*=45; mixed-condition 1, *N*=41, mixed-condition 2, *N*=21. Further, for all subjects in the stimulus-based session and each session of mixed-condition 1, we discarded 38 trials that did not have a pre-test sequence (which were likely surprising and led to poor subject performance), leaving 442 trials for analysis. Also, one subject completed only 432 trials in the rule-based condition, and two subjects completed only 432 trials in mixed-condition 2. Subjects were paid $8 plus a performance bonus per session. The bonus was $1 for each 4% above 50% (max bonus = $12).

Figure 1b shows the timing of each trial. Subjects could make their response from the start of a visual “go cue,” which coincided with the initiation of the test tone, until the trial ended. Subjects were given a 2-s response window, except for 25 subjects in the rule-based condition who were given a 3-s response window. In mixed-condition 2, we did not provide a go cue because the pre-test sequence had a fixed, predictable duration, which served as the go cue. We provided visual and auditory feedback after each trial indicating whether the subject’s response was correct or incorrect. Due to experimenter oversight, the sound played for correct feedback was ∼600 ms longer than the sound played for incorrect feedback. This duration asymmetry could in theory favor incorrect responding. However, given that subjects were monetarily incentivized for correct performance and performed well above chance on average, the duration asymmetry likely had a negligible effect on the results.

## Data analysis

We conducted statistical analyses in R [74] and Matlab. Mixed-effects models were fit using lme4 [75]. When possible, we fit the maximal model (i.e., random intercepts for subjects and random slopes for all within-subjects variables). If the maximal model failed to converge or produced singular fits, we iteratively reduced the random-effects structure until convergence [76]. Post-hoc comparisons were conducted using the emmeans package [77]. Post-hoc multiple comparisons were corrected using the Bonferroni-Holm method [78].

We fit a logistic model to the rule-based and stimulus-based psychometric functions using a maximum-likelihood approach. The logistic model was:

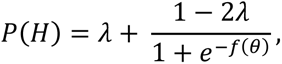

where *P*(*H*) is the probability that the subject chose high frequency, *f*(*θ*) is a linear function predicting choice as a function of task parameters, *θ*, and *λ* is a lapse rate that sets the lower and upper asymptotes of the logistic curve.

For the rule-based condition, choice was predicted as:

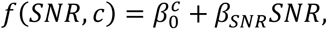

where *β*_*SNR*_ determines the slope of the psychometric function, *SNR* is the signed signal-to-noise ratio of the test tone (positive: high-frequency tones; negative: low-frequency tones), and 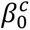 determines the bias (offset). To capture the effects of the rule-based cues, we fit separate 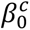 for each prior cue, *c* (low, neutral, or high). We also tested a model in which the slope was free to vary with prior cue.

For the stimulus-based condition, choice was predicted as:

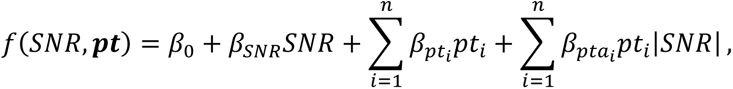

where *pt*_*i*_ is a pre-test tone at position *i* numbered in reverse chronological order (i.e., *pt*_1_ is the tone immediately prior to the test tone), *β*_*pt_i_*_ determines the bias (offset) attributable to each tone, and *β*_0_ is a fixed offset capturing any idiosyncratic biases. The *pt*_*i*_ were contrast coded such that *β*_*pti*_ is the log-odds ratio of choosing high frequency over low frequency at a SNR of 0 (high-frequency regression contrast code: 0.5; low-frequency regression contrast code: −0.5). We modeled an SNR-dependent adaptation-like effect as the interaction term *pt*_*i*_|*SNR*|: |*SNR*| is the unsigned signal-to-noise ratio of the test tone, and each *β*_*ptα_i_*_ determines the influence of that tone on the slope of the psychometric function. At most, we fit data for the last six pre-test tones (i.e., max *n* = 6) because longer tone-burst sequences were infrequent.

To directly compare the effects of the rule- and stimulus-based cues, we fit logistic and linear mixed-effects models to the choice and RT data, respectively. These analyses assessed the three-way interaction between unsigned SNR (lowest [0.05] versus highest [0.5]), the congruence of the cue and the test tone (congruent versus incongruent), and condition (rule-based versus stimulus-based). A significant interaction indicates that the effect of congruence on choice or RT is differentially moderated by SNR between the conditions. For the stimulus-based condition, we included only trials ending with two high or two low pre-test tones (LL and HH trials) to focus on the clearest congruence effects (see Figure 2b). For the choice analysis, the model tests if subjects were more likely to choose correctly/incorrectly when the cue was congruent/incongruent with the test tone and whether this effect varied by SNR and condition. For the RT analysis, the model tests if subjects were faster/slower to respond when the cue was congruent/incongruent. Only correct trials were included in the RT analysis, so congruence between the cue and the test tone is equivalent to congruence between the cue and choice.

### Drift-diffusion modeling (DDM)

We fit DDMs to the data by maximum-likelihood estimation on the full empirical RT distribution, using PyDDM [38]. Optimization was performed using differential evolution [79], a global optimization algorithm that performs well in estimating the parameters of high-dimensional DDMs [38]. Model fits used DDMs in which the two decision bounds corresponded to correct and error choices (as expected by PyDDM). For expositional simplicity, the models are presented with bounds corresponding to high-frequency and low-frequency choices.

In the DDM framework, noisy evidence is accumulated until reaching one of the two bounds, triggering commitment to one of the two choices (in this case, low frequency or high frequency). The DDM was instantiated by the following differential equation:

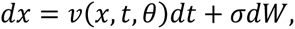

where *x* is the decision variable, *t* is time, *θ* are the parameters determined by the task, *v*(*x*, *t*, *θ*) is the instantaneous rate of evidence accumulation, and *σ* is the instantaneous noise (*σ* = 1). In our implementation, the initial bound height is determined by a parameter *B*, which is equal to half the total distance between the bounds. All models, except as noted, included a linear collapsing bound, which accounts for the truncated RT distributions given that subjects were under time pressure. (RT was the time between go-cue onset and gamepad button press, except for mixed- condition 2 in which a subject’s RT was the time between test-tone offset and button press.) The parameter *t*_*B*_ determines the rate of linear collapse, such that total bound height at time *t* is 2(*B* − *t*_*B*_*t*). Therefore, the DDM process stops, and choice commitment occurs, when |*x*(*t*, *θ*)| ≥ *B* − *t*_*B*_*t*.

The full slope or rate of evidence accumulation was defined as:

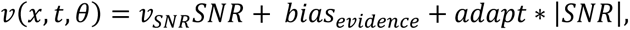

where *v*_*SNR*_ is the drift rate, which influences the rate at which the sensory evidence provided by the test tone in the noisy background, *SNR,* contributes to evidence accumulation.

We defined separate *bias*_*evidence*_ terms for different expectation-generating cue conditions. For the rule-based condition, *bias*_*evidence*_was set to one of three parameters depending on the prior cue (low: *v*_*Low*_; neutral: *v*_0_; high: *v*_*High*_). For the stimulus-based condition, *bias*_*evidence*_ = *v*_0_ + *v*_*Bias*_*pt*_*Bias*_, where *v*_0_ is a fixed offset capturing any idiosyncratic bias toward high or low and *pt*_*Bias*_ is an exponentially weighted sum of the pre-test tones:

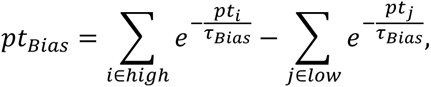

where *pt*_*i*_and *pt*_*j*_ are the sequence positions (numbered in reverse chronological order starting at 0) of the high- and low-frequency tones in the pre-test sequence, respectively. *τ*_*Bias*_is a time constant determining the decay in influence of the tones with increasing temporal distance from the test tone, and *v*_*Bias*_ scales the total influence of the tones on the rate of evidence accumulation.

The adaptation-like effect in the stimulus-based condition was modeled using an SNR-dependent term in the slope, *adapt* ∗ |*SNR*|. The *adapt* quantity was defined as

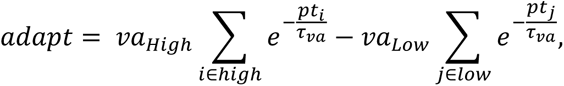

where *τ*_*va*_ is a time constant and *va*_*High*_ and *va*_*Low*_ are separate weights for the high- and low- frequency tones.

Our model also controlled the location between the two bounds at which evidence starts accumulating, *x*_0_. If the starting point is not equidistant between the two bounds, less evidence is required to make one choice versus the other (i.e., a starting-point bias). The starting point was defined as

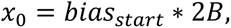

where 2*B* is the total bound height, and *bias*_*start*_ is the bias on the starting point, expressed as a fraction of total bound height [0, 1] (i.e., a value of 0.5 corresponds to an unbiased starting point, equidistant between the bounds). For the rule-based condition, *bias*_*start*_was set to one of three parameters depending on which prior cue was presented on that trial (low: *Z*_*Low*_; neutral: *Z*_0_; high: *Z*_*High*_). For the stimulus-based condition, *bias*_*start*_ = *g*(*Z*_0_ + *Z*_*Bias*_*pt*_*Bias*_). The *pt*_*Bias*_ is the same quantity as in the evidence-accumulation term, *Z*_*Bias*_ scales the total influence of the tones in the pre-test sequence on the starting point, and *Z*_0_ is a fixed offset accounting for idiosyncratic starting-point biases. The function *g(y)* is the logistic function, 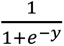, which constrains *bias*_*start*_ between 0 and 1. Although we did not fit the rule-based condition using this logistic transformation, we transformed the resulting parameter estimates to the logit scale to be more comparable with the stimulus-based fits.

In addition to these parameters, all models were fit with a non-decision time, *ndt*_0_, which accounts for the portion of RT that is not determined by decision formation (e.g., sensory or motor processing). For the stimulus-based condition, RTs were faster when the pre-test tones were the same frequency. To account for this effect, total non-decision time was calculated as

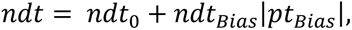

where the extra additive term is the absolute value of the stimulus-based bias calculated above, weighted by *ndt*_*Bias*_. Therefore, the RT for a single instantiation of the DDM process is given by *t*_*s*_ + *ndt*, where *t*_*s*_ is the time at which the bound is reached.

Finally, all models were fit with a lapse rate, *λ*, which mixes the RT distribution predicted by the DDM with a uniform distribution in proportion to *λ* (i.e., *λ* = 0.05 implies that the final predicted distribution is a weighted average of 95% the DDM distribution and 5% a uniform distribution).

We fit models separately to each task condition. The models for the mixed conditions were identical to those described above but included both types of bias. For the rule- and stimulus-based conditions, we also fit reduced models to assess the necessity of different mechanisms for explaining subjects’ behavior. The “base” model was a standard DDM augmented by idiosyncratic bias terms for evidence accumulation and starting point but without cue-dependent biases; it included six parameters, *v*_*SNR*_, *B*, *v*_0_, *Z*_0_, *ndt*_0_, *λ*. The “collapsing-bound” model included one additional parameter, *t*_*B*_. The “full” model in the rule-based condition and the “bias-only” model in the stimulus-based condition included the *bias*_*evidence*_and *bias*_*start*_(and *ndt*_*bias*_for the stimulus-based condition) terms, for a total of 11 parameters each. The “full” model in the stimulus-based condition further included the *adapt* term, for a total of 14 parameters.

For each condition, we also explored the necessity of the different forms of bias by fitting a “starting-point-only” model, which corresponded to the full models with the cue-dependent evidence accumulation bias terms omitted, and an “evidence-only” model, which corresponded to the full models with the cue-dependent starting-point-bias terms omitted. Each of these reduced models had 9 parameters in the rule-based condition and 13 parameters in the stimulus-based condition. Additionally, to test the necessity of the collapsing bound in the presence of the bias terms, we fit a “full, fixed bound” model to the rule-based condition, which was identical to the full model except for having a fixed bound (10 parameters).

Because there is inconsistency across studies in whether RT data are better captured by a collapsing bound or parameter variability [38,80–82], we also fit a model that replaced the *t*_*B*_ parameter in the full rule-based model with drift-rate variability, controlled by a parameter *v*_*width*_ that determines the width of a uniform distribution [83] centered on *v*_*SNR*_. This fixed-bound model with drift-rate variability provided a substantially poorer fit to the data than the full model (mean log-likelihood ratio = −52.90, 0/49 subjects better fit by drift-rate variability model; Figure S1c). We therefore used a collapsing bound in all main analyses.

We used the full-model fits to generate performance functions for each subject. The output of the performance function, *pc*(*Z*, *v*|*c*, *θ*), was the predicted proportion of correct choices for a particular combination of starting-point bias (*z*) and evidence-accumulation bias (*v*), given the expectation- cue context (*c*) and the subject’s other best-fitting DDM parameters (*θ*). For the rule-based condition, the performance function was estimated for the context of the low prior and the high prior. For the stimulus-based condition, the context was set to either two low-frequency pre-test tones (LL) or two high-frequency pre-test tones (HH). These pre-test tones did not change the odds that the test tone would be low or high but nonetheless affected performance, as quantified via bias and adaptation-like effects in the DDM fits.

We estimated the performance function across a grid of *Z*, *v* values (*Z*: (. 1, .9); *v*: [−4,4]), where *Z* dictated the value of *Z*_*Low*_or *Z*_*High*_in the rule-based condition and *Z*_*Bias*_in the stimulus-based condition, and *v* dictated the value of *v*_*Low*_ or *v*_*High*_in the rule-based condition and *v*_*Bias*_in the stimulus-based condition. The predicted proportion correct was generated from the DDM for each point on the grid for all combinations of test-tone frequency and SNR. Next, these values were averaged according to the proportion of each trial type expected in that context. This procedure was implemented separately for each subject, condition, and context. The resulting performance functions were then normalized to a proportion of maximum possible performance scale by dividing by the maximum accuracy obtained for each function. Performance functions were then averaged across subjects for each context, and subject-averaged 97% contours were plotted using the Matlab contour function.

We fit a mixed-effects model to test whether the slope of the relationship between the starting-point and evidence-accumulation biases differed between the rule- and stimulus-based conditions. We entered the starting-point bias as the predictor and assessed the interaction between starting-point bias and bias type (*z_High_, z_Low_*, or *z_Bias_*) in predicting the evidence-accumulation bias. Specifically, we contrasted the slopes of the rule-based and stimulus-based biases ([average of *z_High_* and *z_Low_*] − *z_Bias_*) and the contrast of the rule-based slopes (*z_High_* − *z_Low_*). To confirm that the results were not dependent on which variable was entered as the outcome and which as the predictor, we re-estimated the model with the evidence-accumulation bias as the predictor and the starting-point bias as the outcome, which yielded similar results (rule-based vs. stimulus-based: *B =* 0.47, *t*(53.65) = 2.93, *p =* 0.005; rule-based high vs. low (*B =* −0.21, *t*(118.93) = −0.93, *p* > 0.05). Because the rule- and stimulus-based biases are on different scales, for both models, the bias variables were z*-*scored within condition.

To test the association between rule-based and stimulus-based bias parameters, we calculated the rule-based biases for a given condition as the difference between the high and low bias for both evidence-accumulation (i.e., *v*_*High*_ − *v*_*Low*_) and starting-point (i.e., *Z*_*High*_ − *Z*_*Low*_) biases. We then calculated the Spearman’s correlation between these quantities and *v*_*Bias*_ and *Z*_*Bias*_, respectively.

We generated simulations of the different types of biases for schematic purposes using an analytical solution to the DDM [37]. Unlike some of the fits described above, these simulations had a fixed bound and were not fit to the data but rather used ranges of parameter values that generated psychometric and chronometric functions that were qualitatively consistent with the data.

### Model Comparison

We assessed goodness-of-fit using Akaike information criteria (AIC). We also entered the AIC values into a Bayesian random-effects analysis, which attempts to identify the model among competing alternatives that is most prevalent in the population. This analysis yields a protected exceedance probability for each model, which is the probability that the model is the most frequent in the population, above and beyond chance [84]. Protected exceedance probabilities were computed using the VBA toolbox [85]. We also computed Bayesian information criteria (BIC) values as an additional, more conservative measure of goodness of fit [35]. Finally, to characterize individual differences in DDM fits, we took advantage of the nested nature of our DDM variants and conducted likelihood-ratio tests (LRT) on selected full-versus-reduced DDM comparisons.

### Pupillometry

We recorded each subject’s right-eye position and pupil diameter at a sampling rate of 1000 Hz (EyeLink 1000 Plus; SR Research). Each subject maintained their gaze within a 14.66 x 8.11° window throughout the trial.

If data were missing, either due to blinks or other artifacts, we linearly interpolated the pupil data after removing ±100 ms surrounding the blink (or artifact). We identified additional artifacts by computing the difference between consecutive samples of the pupil time course and removed all high-velocity periods (i.e., > 24 a.u./ms). If the duration of any of these periods exceeded 16 ms, we also removed the ±100 ms of surrounding data. These artifactual periods were then filled via linear interpolation. We also interpolated gaze-position data for time points missing or removed from the pupil time course. We excluded trials in which >50% of the pupil data and sessions in which >60% of the pupil data were missing or were artifactual from further pupil analysis. Additionally, in mixed-condition 1, both sessions had to pass these criteria for the subject to be included in the pupil analysis.

The pupil time course was then low-pass filtered with an 8-Hz cutoff. We z-scored each subject’s time course within each block. For every trial, we calculated the baseline pupil diameter as the average diameter from 0–40 ms relative to test-tone onset, which was then subtracted from the trial time course.

Gaze position for each trial was centered on the average gaze position during the baseline period. For each session, we defined the fixation window as a circle containing 95% of gaze samples across all data (1.36–3.29° visual angle across sessions). To minimize the impact of eye movements on pupil diameter, we excluded trials in which gaze deviated from this window for >15 ms.

Finally, we excluded subjects with fewer than 75 trials remaining in a condition after all cleaning procedures. Overall, we analyzed 44 subjects in the rule-based condition, 42 in the stimulus-based condition, 33 in mixed-condition 1, and 17 in mixed-condition 2.

For statistical analyses, we downsampled the eye data to 50 Hz. We analyzed pupil diameter with respect to the congruence between the cue and choice and restricted our main analyses to correct trials to avoid confounding changes in pupil size with feedback/error related signals or off-task responses. We analyzed the rule- and stimulus-based conditions by fitting a linear mixed-effects model to the data at every time point. These analyses focused on contrasts of the pupil time course as a function of the congruence of choice with the rule- and stimulus-based cues (e.g., incongruent– congruent), and the interaction of congruence with |SNR|. All models controlled for test-tone frequency, |SNR|, baseline pupil diameter, and gaze position. We aligned the data to choice (i.e., the time of the button press).

To test the association between behavioral bias and the congruence effects in the rule-based condition, separate linear models were fit to the pupil data per subject to obtain estimates of individual-subject congruence contrasts. These contrasts were then correlated across time with the estimate of behavioral bias obtained from the logistic fits to the psychometric function described above, where bias was defined as 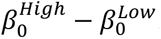. We used FDR correction to adjust *p*-values for multiple comparison across time [86, 87]. Congruence contrasts were corrected together across time and contrast.

## Acknowledgements

YEC, JIG, and LSA received support for this work from an Office of Naval Research grant [N000141612539]. NT was supported by a T32 training grant from the National Institutes of Health [MH014654].

## Data availability

The datasets generated and analyzed for this article are available at: https://osf.io/f9nyr/. The analysis code for this article is available at: https://github.com/TheGoldLab/Analysis_Tardiff_etal_AuditoryPriors.

