## Supplemental Figures 1 and 2 for "Rule-based and stimulus-based cues bias auditory decisions via different computational and physiological mechanisms"

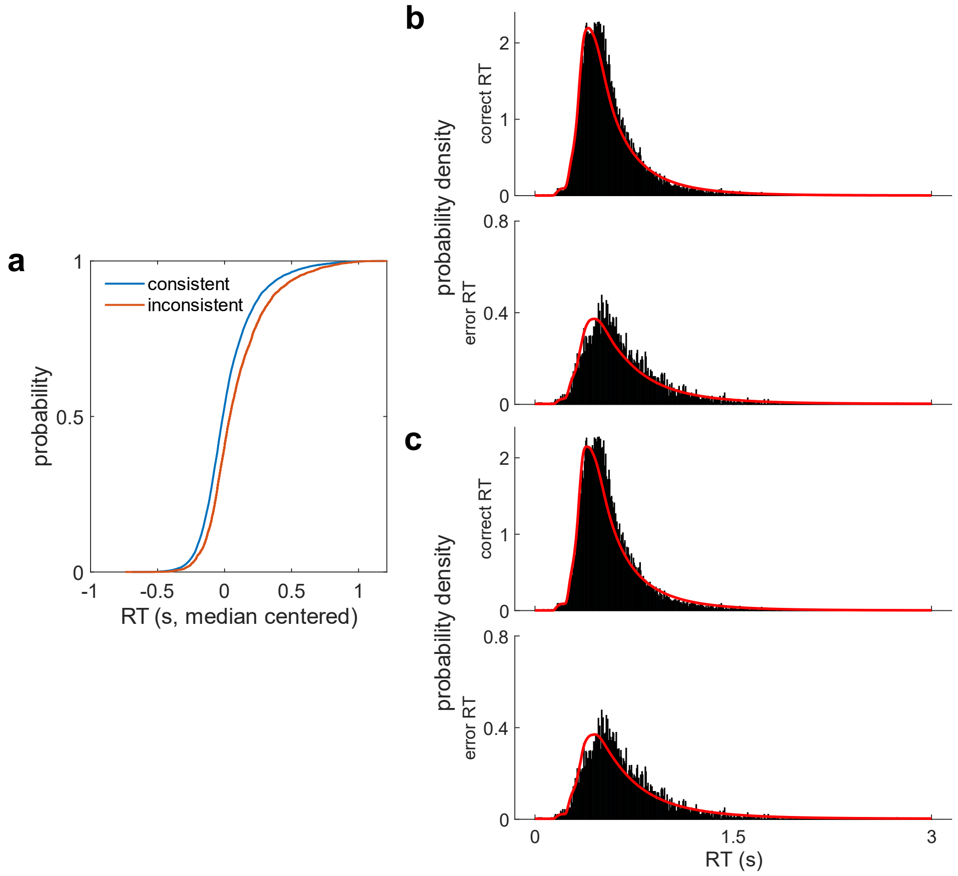


**Figure S1**. **RT distributions and alternative-bound models. a** Empirical cumulative density function for the stimulus-based condition RTs, plotted separately for trials in which the last two pre-test tones were consistent (i.e., HH, LL) or inconsistent (i.e. HL, LH). **b** Fits of the full, fixed bound model to RT data, averaged across subjects. **c** Fits of the drift-variability model to RT data, averaged across subjects. For **a**, RT data were median-centered within subjects to remove between-subjects variance in average RTs and then pooled across subjects. In **b** and **c**, red lines are the probability density functions (PDFs) from each model. Bars are the data RT histograms, normalized to yield an empirical PDF. By convention, correct RTs and error RTs jointly integrate to one, so differences in total density between correct and error RT distributions reflect differences in the rate of correct and error responses.

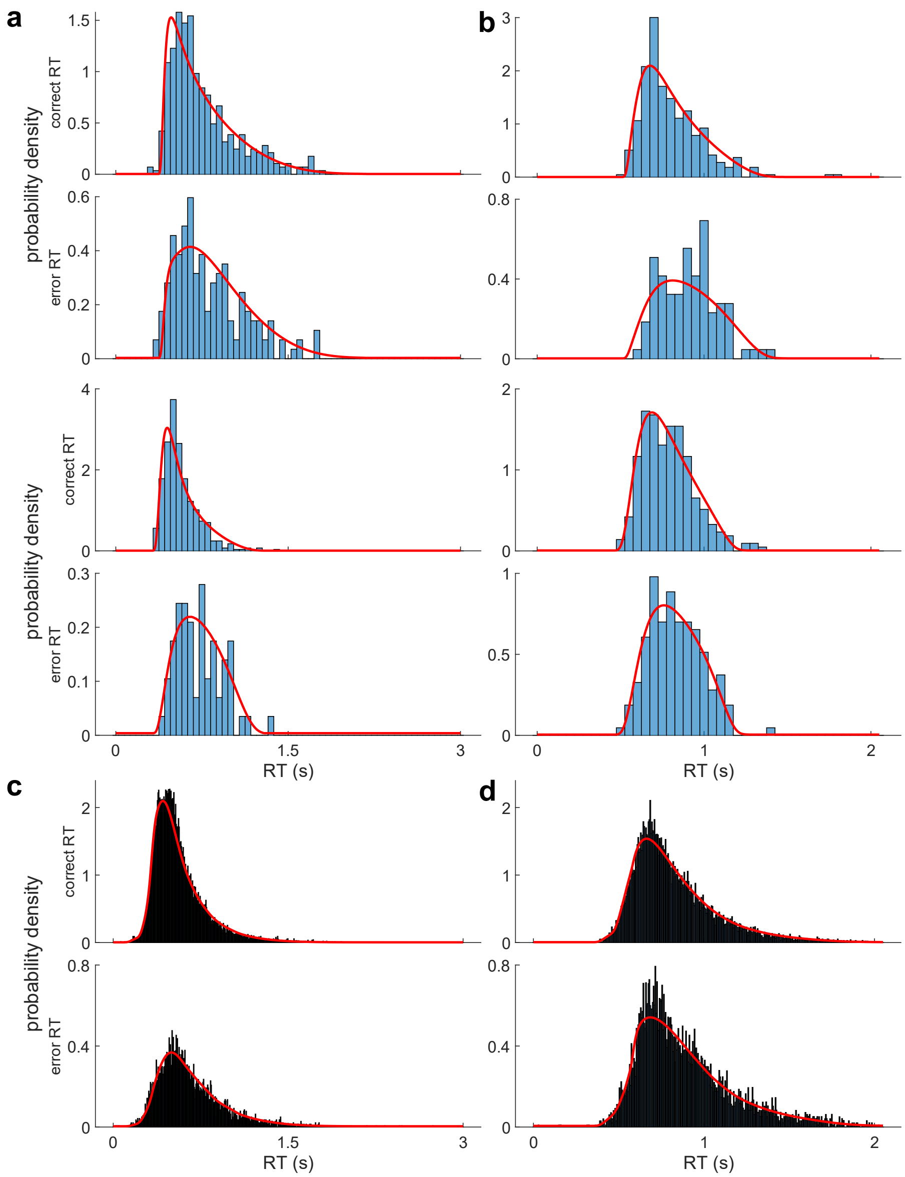


**Figure S2**. **Best DDM fits to RT distributions**. **a** Fits of the full rule-based DDM to two example subjects' RT data. **b** Fits of the full stimulus-based DDM to two example subjects' RT data. **c, d** Same as **a, b**, averaged across subjects. Empirical and model PDFs were plotted identically to the PDF plots in S1.
